# Cowpea Chlorotic Mottle Virus virus-like particles as DNA or RNA delivery vehicle to fish cells

**DOI:** 10.64898/2026.10.15.759507

**Authors:** Mark R. G. Goldman, Shuqin Cao, Jeroen J. L. M. Cornelissen, Richard Kormelink, Maria Forlenza

## Abstract

Cowpea Chlorotic Mottle Virus (CCMV) is an icosahedral plant virus composed of 180 monomeric units of the capsid protein. Virions encapsulate the viral RNA genome, which can be removed from the virion by changing pH conditions, after which the capsid proteins can be reversibly assembled to form empty CCMV virus-like particles (VLPs), or VLPs that have encapsulated either heterologous DNA or RNA. Earlier research has shown that CCMV VLPs are suitable as delivery vehicles for heterologous DNA/RNA molecules into mammalian cells, leading to subsequent expression of their encoded genes. In this study, we evaluated the potential of CCMV VLPs to encapsulate nucleic acids and transfer them to fish cells *in vitro,* and to zebrafish larvae and juveniles *in vivo*. VLPs containing pVax-GFP plasmid (3690 bp) were stable for up to 5 weeks in buffer at pH 7.2 and pH 5, whereas VLPs containing GFP-mRNA (1168 nt) were stable for up to 3 weeks, but only when stored in a buffer at pH 4.5. When administered directly to the culture medium, VLPs were readily taken up by fish as well as mammalian cell lines but could not unpack and transfer nucleic acids to the cells. VLPs delivered by transfection however, could transfer nucleic acids to the cells, leading to reporter gene expression. When VLPs were administered directly to the culture medium along with the cell-penetrating peptide dfTAT, only VLPs containing pVax-GFP in pH 7.2 buffer were able to transfer DNA to mammalian but not fish cells. *In vivo* however, upon intramuscular injection, the same VLPs were able to transfer plasmid DNA to myocytes of zebrafish larvae and juveniles as efficiently as when injecting naked plasmid DNA. Altogether, these findings provide the foundation for future studies on the use of CCMV VLPs as delivery vehicles for DNA or RNA to fish cells.

## Introduction

Cowpea chlorotic mottle virus (CCMV) is a small icosahedral plant virus, known to infect cowpea plants (*Vigna unguiculata)* [1]. In its native form, CCMV virions consist of 180 monomeric units of the capsid protein (CP) [2]. Virions encapsulate the viral RNA genome, which can be removed from the core by (partial) virion disassembly upon changing pH conditions, after which the CP can be reversibly assembled to form empty CCMV virus-like particles (VLPs) [3]. This property has not only been used to exchange CCMV RNA with other heterologous RNA [4–7], but also to exchange it with DNA [7–10], fluorescent proteins [11,12], synthetic polymers [13–15], enzymes [16,17] and gold nanoparticles [18,19]. Earlier research showed that the size and shape of CCMV VLPs depend on their cargo. Single-stranded RNA molecules shorter than 3200 nucleotides are encapsulated within a single spherical VLP, whereas longer RNA molecules are encapsulated in dimers, trimers, or multiplets VLPs that share the long RNA molecule [20]. In contrast, double-stranded (ds)DNA molecules longer than 100 base pairs are encapsulated in rod-shaped VLPs, whereas only short dsDNA molecules (≤ 50 base pairs) are encapsulated in spherical VLPs [7]. While the potential of CCMV VLPs as scaffold and cargo carrier has been explored, contradictory results are reported on their suitability to deliver nucleic acids to cells *in vitro*. VLPs containing *eGFP-mRNA* are able to transfer the mRNA to HEK293, HeLa and HK2 cells when directly added to the culture medium, leading to eGFP expression levels similar to those from cells that received *eGFP-mRNA* by transfection [6]. In contrast, mammalian BHK-21 cells showed fluorescence only when receiving VLPs by transfection using Lipofectamine 2000 [21]. To overcome the use of transfection agents, Lam & Steinmetz administered VLPs in combination with the cell-penetrating peptide M-lycotoxin peptide L17E [22], known to disrupt endolysosomal membranes and promote uptake by micropinocytosis [23]. They succeeded in silencing 30% of GFP expression in HeLa cells by administering VLPs containing GFP-specific siRNAs. The addition of the L17E peptide led to a further reduction to about 50% [22]. A similar reduction in FOXA1 expression levels was obtained in MCF-7 cells upon administration of L17E-conjugated VLPs containing FOXA1-specific siRNAs, whereas no downregulation occurred in the absence of the cell-penetrating peptide [22]. *In vivo*, VLPs encapsulating immunostimulatory oligodeoxynucleotides (ODNs) have been used in colon cancer and melanoma mouse models, and were shown to enhance the efficacy of ODN1826 compared to the naked ODN1826, ultimately leading to prolonged survival and slower tumour growth [8]. Furthermore, oral and intramuscular administration of VLPs containing white spot syndrome virus (WSSV) glycoprotein VP28 gene-specific siRNAs successfully protected shrimps against a challenge with WSSV, leading to up to 40% protection after oral administration, and up to 100% protection after intramuscular administration [24].

DNA vaccines are notoriously effective in fish, especially in protection against viral diseases [25–29]. The first DNA vaccine, APEX-IHN®, was licensed in Canada in 2005 and it is still in use to protect Atlantic salmon against Infectious Hematopoietic Necrosis Virus (IHNV) [30]. More recently, also RNA vaccines were shown to be effective in fish. Lipid nanoparticle-mRNA (LNP-mRNA) complexes have successfully been used to deliver eGFP-mRNA to zebrafish and Atlantic Salmon [31,32]. Furthermore, vaccination with LNP-mRNA encoding the glycoprotein of viral haemorrhagic septicaemia virus (VHSV) in rainbow trout was able to induce neutralizing antibodies and protect rainbow trout against a lethal challenge with VHSV [32]. Encapsulation methods for nucleic acid vaccines must not only protect the nucleic acid cargo against degradation but also transfer it to host cells. Currently, encapsulation methods used in fish include alginate, chitosan, liposomes, MicroMatrix^TM^ and Poly D,L-lactic-co-glycolic acid (PLGA) [29,33,34].

In this study, taking advantage of the ease of production and high yields obtained when CCMV is produced in cowpea plants, the straightforward assembly of CCMV VLPs with nucleic acids, and the reported ability to deliver nucleic acids at least to mammalian cells, we investigated the potential of CCMV VLPs to encapsulate nucleic acids for delivery to fish cells *in vitro,* and to zebrafish larvae and juveniles *in vivo*. To this end, rod-shaped VLPs containing pVax-GFP and spherical VLPs containing *GFP-mRNA* were assembled and assessed for their stability and efficacy of nucleic acid delivery into fish cells. Furthermore, we investigated two strategies to facilitate and improve the release of nucleic acids in fish cells *in vitro*: (1) inhibition of endosomal acidification using chloroquine, and (2) endosomal escape using the cell-penetrating peptide dfTAT (dimer fluorescent TAT). Results from this study provide evidence of the potential of CCMV VLPs for delivery of nucleic acid-based vaccine to fish.

## Materials and methods

### Nucleic acids

*GFP*, from in-house pcDNA3-GFP (10 µg) [35] and pVax (2 µg) (Invitrogen) were restricted using *EcoRI* (Promega, Cat. no. R601A) and *XbaI* (Promega, Cat. no. R618G). The products were gel-purified using the QIAquick gel extraction kit (Qiagen, Cat. no. 28704), after which *GFP* was ligated into pVax using T4 ligase (Promega, Cat. no. M1794) to generate pVAX-GFP. The plasmid was transformed into JM109 competent cells (Promega, Cat. no. L2001), after which the transformed bacteria were propagated in LB medium, pVax-GFP was isolated using the EndoFree Plasmid Giga Kit (Qiagen, Cat. no. 12391).

*GFP-mRNA* was generated using 1 µg PCR product as a template. The PCR product was produced using pVAX-GFP as template and by amplifying a 1018 bp product that includes the T7 promotor and the *GFP* gene using the pVax-GFP_forward: CCACTGCTTACTGGCTTATC and pVax-GFP_reverse: AGAATGACACCTACTCAGAC primers. Plasmid DNA was removed using *DpnI* (New England Biolabs, Cat. no. R0176S), after which the PCR product was purified using the DNA Clean & Concentrator-25 kit (Zymo Research, Cat. No. D4033). *GFP-mRNA* was generated using the HiScribe® T7 ARCA mRNA Kit, with tailing (New England Biolabs, Cat. no. E2060S), after which *GFP-mRNA* was purified using the Monarch® RNA Cleanup Kit (New England Biolabs, Cat. no. T2040L). The size and quality of the RNA was assessed on an agarose gel and *GFP-mRNA* was stored at -80°C.

### CCMV purification from cowpea plants

CCMV was purified from infected cowpea plants according to the protocol described by Bancroft [36], with minor modifications. Briefly, 14-day-old cowpea plants were infected with CCMV by gently rubbing virus inoculum (prepared by grinding 3-4 virus infected leaves in 10 mL water) on the first two leaves with silicon carbide (400 mesh; Sigma, Cat. no. 357391). After 10-14 days, leaves were harvested and blended in cold homogenization buffer (2 mL per gram of tissue; 200 mM Na-acetate (Merck, Cat. no. 106268), 10 mM ascorbic acid (Sigma, Cat. no. A92902), 10 mM EDTA (Merck, Cat. no. 108418), pH 4.8). The plant extract was filtered through a double layer of cheesecloth to remove large plant particles and left at 4°C for an hour to allow plant proteins to precipitate, prior to centrifugation at 1000 rpm for 10 minutes at 4°C. The supernatant was collected and filtered through a funnel containing glass wool to filter the remaining plant material. Next, 10% (w/v) polyethylene glycol 6000 (Merck, Cat. no. 807491) was added to precipitate virions during 1 hour at 4°C and under gentle agitation at 300 rpm. Virus was pelleted by centrifugation at 10.000 rpm for 15 minutes at 4°C, after which pellets were resuspended in cold virus storage buffer (100 mM Na-acetate, 1 mM EDTA, 1 mM Na-azide, pH 5.0). The virus was further purified on a 37.5% (w/w) cesium chloride density gradient (Sigma, Cat. no. C-3032), during centrifugation at 40.000 rpm for 16 hours at 10°C in a Beckman Coulter Optima LE-80K ultracentrifuge with a SW 55 Ti swinging-bucket rotor. Bands containing purified virus particles were collected and dialyzed three times in a 12-14 kDa molecular weight cut-off (MWCO) membrane (Spectrum, Cat. no. 132754) against cold virus storage buffer and stored at 4°C until further use.

### CCMV labelling

CCMV was fluorescently labelled according to the protocol optimised by de Ruiter [7]. This protocol was optimized to have a degree of labelling of 0.05 dyes per capsid protein, and gives maximum fluorescence with minimum hindrance during assembly of VLPs. In brief, CCMV was dialysed three times against PBS (supplemented with 10 mM MgCl2, pH 7.5) in a 12-14 kDa MWCO dialysis membrane at 4°C, after which the virus concentration was measured, and for further application was diluted to ± 2 mg/mL. ATTO 647N-NHS (ATTO-TEC, Cat. no. AD647N-31) was dissolved in 200 µL DMSO (Merck, Cat. no. 102952), after which 0.003% (w/w) ATTO 647N-NHS was mixed with CCMV and incubated for 1 hour on a roller mixer at 21°C. Unbound dye was removed by dialysing three times at 4°C against PBS in a 12-14 kDa MWCO dialysis membrane, after which the labelled virus (referred to as ATTO647N-CCMV) was dialysed another three times against virus storage buffer at 4°C. Virus was stored at 4°C (for up to 2 months).

### Isolation and purification of CCMV Capsid protein

CCMV or ATTO647N-CCMV were dialyzed in a 6-8 kDa MWCO dialysis membrane (Spectrum, Cat. no. 132650) overnight at 4°C against a 1000 times excess volume of RNA precipitation buffer (50 mM Tris-HCl, 1 M CaCl2, pH 7.5). The next day, precipitated RNA was removed by centrifugation for 2 hours at 40.000 rpm in a Beckman Coulter Optima LE-80K ultracentrifuge in a SW 55 Ti swinging-bucket rotor. The supernatant containing the soluble CP dimers was collected and next dialysed three times against 5x assembly buffer (250 mM Tris-HCl, 50 mM KCl, 500 mM NaCl, 25 mM MgCl2, pH 7.2) in a 3.5 kDa MWCO dialysis membrane at 4°C. Soluble CP dimers were diluted in 5x assembly buffer to ± 1-2 mg/mL and stored for a maximum of 2 weeks at 4°C.

### Encapsulation of nucleic acids into CCMV VLPs

Nucleic acids were diluted in nuclease-free water and added dropwise to CPs (in 5x assembly buffer, diluting the mixture to a final concentration of 1x assembly buffer (50 mM Tris-HCl, 10 mM KCl, 100 mM NaCl, 5 mM MgCl2, pH 7.2). The mixture was incubated overnight at 4°C while rotating at 20 rpm. First, we optimised the CP:DNA (w/w) ratio that could encapsulate all pVax-GFP by mixing CPs and pVax-GFP at different ratios in 1x assembly buffer. After overnight incubation at 4°C while rotating at 20 rpm, products were analysed on an agarose gel.

VLPs assembled with pVax-GFP (referred to as VLPs-DNA) were obtained using a 6:1 (CP:DNA (w/w)) ratio, after which part of the VLPs-DNA were dialysed overnight from assembly buffer at pH 7.2 to VLP-DNA buffer at pH 5.0 (50 mM Na-acetate, 1 M NaCl and 1 mM Na-azide, pH 5.0) in a 12-14 kDa MWCO membrane at 4°C.

VLPs containing *GFP-mRNA* (referred to as VLPs-RNA) were obtained by mixing CP and RNA in a 6:1 (w/w) ratio, after which part of the VLPs-RNA were dialysed overnight from assembly buffer at pH 7.2 to VLP-RNA buffer at pH 4.5 (50 mM Na-acetate, 8 mM Mg-acetate, 1 mM Na-azide, pH 4.5) in a 12-14 kDa MWCO membrane at 4°C; part of the obtained VLPs-RNA at pH 4.5 were dialysed again against 1x assembly buffer at pH 7.2 overnight at 4°C.

For long-term storage, VLPs-DNA were either stored in assembly buffer at pH 7.2 or in VLP-DNA buffer pH 5.0, whereas VLPs-RNA were stored in VLP-RNA buffer pH 4.5; all VLPs were stored for a maximum of 4 weeks at 4°C.

### Nuclease treatment

To assess whether VLPs were able to protected the encapsulated nucleic acids, a nuclease treatment was performed. To this end VLPs-DNA were mixed with 1 µL DNase solution (2.5 µL Benzonase® (25-29 U/uL, Merck, Cat. no. 70746), 19 µL nuclease-free water, 10 µL 0.5 M MgCl2); VLPs-RNA were mixed with 1 µL of 2 µg/mL RNase A (Roche, Cat. no. 10109142001), and next incubated for 15 minutes at 21°C. In parallel, pVax-GFP and *GFP-mRNA* were used as positive controls to assess successful nuclease treatment. Considering that VLPs-DNA were stored in buffer at pH 5.0 and VLPs-RNA in buffer at pH 4.5, an additional control was added in which pVax-GFP and *GFP-mRNA* were resuspended in the same pH buffer as the corresponding VLPs.

### Transmission electron microscopy

VLPs (10 μL) were applied onto glow-discharged (VLPs-RNA only) formvar-carbon coated grids (Electron Microscopy Sciences, Cat. no. FCF200-CU) for 5 minutes, after which excess liquid was removed using filter paper. Grids were negatively stained with 2% (w/v) uranyl acetate (Electron Microscopy Sciences, Cat. no. 22400); after 30 seconds, excess liquid was drained using filter paper, and grids were dried for a minimum of 60 minutes at 21°C. Images of VLPs were made on the JEM-1400Plus Transmission Electron Microscope (JEOL).

### Cells

*Epithelioma papulosum cyprini* (EPC) cells were cultured in Leibovitz’s L-15 Medium (L15) (Gibco, Cat. no. 11415-049), supplemented with 10% Foetal Bovine Serum (FBS) (Gibco, 26140-079) and 100 units/mL penicillin - 100 μg/mL streptomycin (Gibco, Cat. no. 15140-122) at 27°C. *Henrietta Lacks* (HeLa) cells were cultured in Minimum Essential Medium (MEM) with 2 mM L-glutamine and 25 mM HEPES (Gibco, Cat. no. 31095-029), supplemented with 10% FBS and 100 units/mL penicillin - 100 μg/mL streptomycin at 37°C in the presence of 5% CO_2_.

### Fluorescent microscopy of VLP uptake and GFP reporter gene expression

To evaluate VLP uptake in EPC and HeLa cells, cells were seeded in 24-well plates (100.000 cells/well) and incubated overnight at 27°C (EPC cells) or 37°C with 5% CO_2_ (HeLa cells). The next day, ATTO647N-VLPs-DNA pH 7.2 equivalent to 0.8 µg pVax-GFP and ATTO647N-VLPs-RNA pH 7.2-4.5 equivalent to 0.5 µg GFP-mRNA were added and cells were cultured for 4 hours at 27°C (EPC cells) or 37°C with 5% CO_2_ (HeLa cells). After 4 hours, cells were trypsinized, centrifuged at 400 x *g* for 5 minutes, and pellets were resuspended in complete culture medium. Cells (100 µL) were then transferred onto a Poly-L-Lysine-coated microscopy slide and incubated for 1 hour at 27°C (EPC cells) or 37°C (HeLa cells). The following incubation and washing steps were carried out at 21°C; cells were fixed with 4% paraformaldehyde (PFA; Merck, Cat. no. 818715) for 15 minutes, washed twice with PBS for 5 minutes, and nuclei were stained with DAPI (Sigma, Cat. no. D9542-5MG) for 10 minutes. After two washes with PBS for 5 minutes each, cell surface was stained with 5 µg/mL Alexa Fluor 488 Wheat Germ Agglutinin (WGA-488, Invitrogen, Cat. no. W11261) in PBS for 20 minutes, after which the cells were washed twice with PBS for 5 minutes. Cells were kept in PBS at 4°C and imaged within 24 hours. Cells were imaged using the Nikon Ti Eclipse PFS3 microscope in combination with the Andor Revolution spinning disk confocal (Yokogawa). Cells were imaged using the 60x oil objective (Nikon Plan Apo VC; 1.4 oil NA; MRD01602) and the 100x oil objective (Nikon Plan Apo VC; 1.4 oil NA; MRD01901) with the following settings: DAPI excitation 405 nm, emission 417-477 nm, exposure 600 ms, EM gain 1000, digitizer 10 MHz (14-bit); GFP excitation 488 nm, emission 510-540 nm, exposure 600 ms, EM gain 1000, digitizer 10 MHz (14-bit); ATTO647N excitation 650 nm, emission 665-705 nm, exposure 800 ms, EM gain 1000, digitizer 10 MHz (14-bit); BF DIC, exposure 400 ms, EM gain 300, digitizer 10 MHz (14-bit).

To assess GFP expression in cells to which VLPs were either directly added to the cell culture medium or transfected into the cells, cells treated with chloroquine, or cells treated with dfTAT, samples were processed according to the following protocol. Cells to which VLPs-DNA were added to the cell culture medium or cells that were transfected with pVax-GFP or VLPs-DNA were washed with PBS, fixed with 4% PFA for 15 minutes, washed with PBS for 5 minutes, and nuclei were stained with DAPI for 10 minutes. All steps were performed at 21°C. Cells were kept in PBS at 4°C and imaged within 24 hours. Cells to which VLPs-RNA were either directly added to the cell culture medium or cells that were transfected with *GFP-mRNA* or VLPs-RNA were washed with PBS, fixed with 4% PFA for 15 minutes, and permeabilized with 0.1% triton X-100 for 15 minutes. After one wash with PBS, cells were incubated with 1 µg/mL Chicken (IgY) polyclonal anti-GFP (Antibodies incorporated, Cat. no. GFP-1020, AB_2307313), washed 3 times with PBS, and then incubated with 2 µg/mL Goat Anti-Chicken IgY Alexa Fluor-488 (Abcam, Cat. no. ab150173, Abcam, AB_2827653). After three washes with PBS, nuclei were stained with DAPI for 10 minutes. All steps were performed at 21°C. Cells were kept in PBS at 4°C and imaged within 24 hours using a DMi8 fluorescence microscope (Leica Microsystems), controlled by Leica LASX software (version 3.6.0), equipped with a 10x (HC PL FLUOTAR 10x/0.32 DRY, Cat. no. 11506522) and 20x (HC PL FLUOTAR L 20x/0.40 DRY, Cat. no. 11506243) objectives and DFC3000 G camera (Leica Microsystems). Image acquisition was performed using the following settings: BF excitation, exposure 3-8 ms, gain 1; DAPI excitation, exposure 80-100 ms, gain 1; GFP excitation, 200 ms, gain 1; dfTAT (5-Carboxytetramethylrhodamine) excitation, exposure 150 ms, gain 1. Only for cells treated with VLPs in combination with dfTAT, a different GFP exposure time was used: 500 ms.

### Transfection of nucleic acids and VLPs

EPC and HeLa cells were seeded in 24-well plates (100.000 cells/well) and incubated overnight at 27°C (EPC cells) or 37°C with 5% CO_2_ (HeLa cells). The next day, VLPs-DNA, equivalent to 0.8 µg pVax-GFP, or VLPs-RNA, equivalent to 0.5 µg *GFP-mRNA*, which were nuclease-treated (or not), were either directly added to the cells in the cell culture medium or were transfected into the cells. For transfections of VLPs, VLPs-DNA were diluted in 25 µL nuclease-free water, and 2.8 µL FuGENE® HD Transfection Reagent (Promega, Cat. no. E2311) was added. After 10 minutes at 21°C, the transfection mixture was added to the cells; VLPs-RNA were diluted in 25 µL Opti-MEM (Gibco, Cat. no. 31985062), and 1.5 µL Lipofectamine 2000 (Invitrogen, Cat. no. 52758) was added. After 5 minutes at 21°C, the transfection mixture was added to the cells. In parallel, 0.8 µg pVax-GFP or 0.5 µg *GFP-mRNA* was used as positive control for DNA or RNA transfection, respectively. Considering that VLPs-DNA were stored in VLP-DNA buffer at pH 5.0 and VLPs-RNA in VLP-RNA buffer at pH 4.5, as an additional transfection control, 0.8 μg pVax-GFP or 0.5 μg *GFP-mRNA* was resuspended in the same buffer and in an equivalent volume (25 μl) used for the VLPs, before adding the transfection reagent. Transfected cells were incubated for 5 hours, after which the medium was replaced with fresh culture medium, and were cultured for an additional 3 days. In cells to which the VLPs were added directly to the cell culture medium, the VLPs were kept on the cells for 3 days. After 3 days, samples were treated and imaged as described above and GFP fluorescence was quantified by flow cytometry as described below.

### Inhibition of endosomal acidification using chloroquine

A 0.1 M chloroquine (Sigma, Cat. no. C6628) stock solution was made by diluting chloroquine in Minimum Essential Medium (MEM) supplemented with 2 mM L-glutamine and 25 mM HEPES, at pH 7.5, after which the solution was filter sterilised, aliquoted and stored at -20°C. To evaluate chloroquine toxicity, EPC and HeLa cells (25.000 cells) were seeded in triplicate wells of a 96-well plate and incubated with a 4x serial dilution of chloroquine (0.5 µM to 1230.2 µM) in culture medium. After 24 hours at 27°C (EPC cells) or 37°C with 5% CO_2_ (HeLa cells), plates were fixed with 4% PFA for 60 minutes, and stained with 1% crystal violet (Merck, Cat. no. 1.01408) for 30 minutes. The OD was measured at 595 nm (Multiskan GO, Thermo Scientific), and cell viability was calculated relative to unstimulated control cells.

Selected concentrations were then used in combination with VLPs. EPC and HeLa cells were seeded in 24-well plates (100.000 cells/well) and treated with chloroquine (2, 10, or 50 µM) for 10 minutes at 27°C (EPC cells) or 37°C with 5% CO_2_ (HeLa cells). Next, VLPs-DNA equivalent to 0.8 µg pVax-GFP and VLPs-RNA equivalent to 0.5 µg *GFP-mRNA* were added to the cells, and cells were cultured for 3 days. After 3 days, samples were treated and imaged as described above and GFP fluorescence was quantified by flow cytometry as described below.

### Administration of cell-penetrating peptide dfTAT

A 1 mM stock solution was made by dissolving dfTAT (MedChemExpress, Cat no. HY-P3422-5; Cambridge Research, Cat no. crb1001690j) in nuclease-free water, after which the solution was filter sterilised, aliquoted and stored at -80°C.

To assess dfTAT uptake in EPC and HeLa cells in combination (or not) with VLPs-DNA, EPC and HeLa cells were seeded in 24-well plates (100.000 cells/well) and incubated overnight at 27°C (EPC cells) or 37°C with 5% CO_2_ (HeLa cells). The next day, cells were incubated for 0.25-5 hours with 5 µM dfTAT in combination (or not) with VLPs-DNA, equivalent to 0.8 µg pVax-GFP, after which the cells were washed with PBS, trypsinized and measured by flow cytometry as described below.

To investigate whether VLPs were able to release the nucleic acids to the cytoplasm of EPC and HeLa cells when administered in combination with dfTAT, EPC and HeLa cells were seeded in 24-well plates (100.000 cells/well) and incubated overnight at 27°C (EPC cells) or 37°C with 5% CO_2_ (HeLa cells). The next day, cells were incubated for 5 hours with 1-40 µM dfTAT in combination (or not) with VLPs-DNA, equivalent to 0.8 µg pVax-GFP or VLPs-RNA, equivalent to 0.5 µg *GFP-mRNA*, after which the medium was replaced with fresh culture medium. Cells were then incubated for 2-3 days at 27°C (EPC cells) or 37°C (HeLa cells). After 2-3 days, samples were treated and imaged as described above and GFP fluorescence was quantified by flow cytometry as described below.

### Flow cytometry

For cells to which the dfTAT, pVax-GFP or VLPs-DNA were directly added to the culture medium, or for cells that were transfected with pVax-GFP or VLPs-DNA, cells were washed with PBS and trypsinized with 0.05% trypsin-EDTA (Gibco, Cat. no. 25300-104; EPC cells) or 0.25% trypsin-EDTA (Gibco, Cat. no. 25200-072; HeLa cells), after which the same volume FACS buffer (RPMI) (Gibco, Cat. no. 52400-025) supplemented with 0.5% BSA (Roche Diagnostics GmbH, Cat. no. 10735086001) and 0.01% sodium azide (Sigma, Cat. no. 8.22335.0100) was added. For cells to which the *GFP-mRNA* or VLPs-RNA were directly added to the culture medium, or for cells that were transfected with *GFP-mRNA* or VLPs-RNA, cells were washed with PBS and trypsinized with 0.05% trypsin-EDTA (EPC cells) or 0.25% trypsin-EDTA (HeLa cells). Cells were centrifuged at 400 x g for 5 minutes, and pellets were resuspended in BD Cytofix/Cytoperm (BD, Cat. no. 554715) and incubated for 20 minutes. After two washes with BD Perm/Wash buffer (BD, Cat. no. 554715), cells were incubated with 1 µg/mL Chicken (IgY) polyclonal anti-GFP in FACS buffer, washed 3 times with FACS buffer, and then incubated with 2 µg/mL Goat Anti-Chicken IgY Alexa Fluor-488 in FACS buffer. After three washes with FACS buffer, cells were resuspended in FACS buffer and at least 10.000 events were measured on CytoFLEX LX (Beckman Coulter, USA). Data were analysed using FlowJo (FlowJo LLC, v10.10.0).

### Zebrafish lines and maintenance

Adult and juvenile zebrafish (AB strain) were reared in the Aquatic Research Facility (ARF) of Wageningen University and Research. Adult zebrafish were fed commercial dry feed (Caviar 300-500 μm, BernAqua) twice daily at 2.5% body weight per day. Juveniles (5-week-old) were fed artemia and dry feed (Caviar 100-200 μm, BernAqua) at 2.5% body weight per day. Zebrafish were kept according to local animal welfare regulations of The Netherlands, and handled according to standard protocols [37,38]. Zebrafish larvae were reared in E3 medium (0.44 mM CaCl_2_, 0.18 mM KCl, 0.40 mM MgCl_2_·6H20, 4.96 mM NaCl in demi water at pH 7.2) at 28°C. During all experiments, the health status and welfare of zebrafish were monitored visually daily. No abnormal behaviour and/or health problems were observed. Therefore, no humane endpoints were applied during the experiments.

The following lines were used in this study: optically transparent casper [39], transgenic *Tg(mpeg1:eGFP)^gl22^* marking macrophages with GFP under the control of *mpeg1* (Macrophage expressed gene-1) promotor [40], and *Tg(mpx:eGFP)^i114^* marking neutrophils with GFP under the control of *mpx* (myeloperoxidase) promotor [41]. Competences of all researchers and biotechnician involved in animal experiments, and all animal experiments with zebrafish older than 5 days post fertilization (dpf) were approved by the local experimental animal committee (project number 2023.W-0019.001).

### Intramuscular injection

Prior to injection, zebrafish were anaesthetized in 0.017% Ethyl 3-aminobenzoate methanesulfonate (MS-222) (Sigma, Cat. no. A5040) in E3 medium. To assess VLP uptake by macrophages or neutrophils, 3 dpf *Tg(mpeg1:eGFP)* and *Tg(mpx:eGFP)* zebrafish larvae (n=5 per group) were injected intramuscularly with 2 nl of assembly buffer pH 7.2 (as control), or 2 nl of 0.207 mg/mL ATTO647N-VLPs-DNA pH 7.2. After injection, zebrafish were transferred to fresh E3 medium and imaged 3 to 4 hours post-injection as described below. To assess whether VLPs could transfer DNA to zebrafish larvae, 3 dpf casper zebrafish larvae (n=35 per group) were injected intramuscularly with 2 nl of 0.207 mg/mL DNase treated pVax-GFP (as control), 2 nl of 0.207 mg/mL pVax-GFP and 2 nl of DNase-treated 0.207 mg/mL ATTO647N-VLPs-DNA pH 7.2. After injection, zebrafish were transferred to fresh E3 medium and imaged 5 days post-injection, as described below. To assess whether VLPs could transfer DNA to zebrafish juveniles, 5-week-old casper zebrafish juveniles (n=10 per group) were either left unhandled or injected intramuscularly with 1-2 µL of 0.221 mg/mL pVax-GFP or 1-2 µL of DNase-treated 0.221 mg/mL ATTO647N-VLPs-DNA pH 7.2. After injection, zebrafish were transferred to fresh tank water and imaged 8 days post-injection, as described below.

### *In vivo* imaging of zebrafish

Prior to imaging of zebrafish larvae, individuals were anaesthetized in 0.017% MS-222, after which they were embedded in 1 to 1.2% UltraPureTM Low Melting Point Agarose (Thermo Fisher, Cat. no. 16520-050) at the bottom of a 35 mm petri dish with a 14 mm uncoated glass No. 0 coverslip (MatTek corporation, Cat. no. P35G-0-14-C). Images were acquired using a Nikon Ti Eclipse PFS3 microscope in combination with the Andor Revolution spinning disk confocal (Yokogawa), using the 20x objective (Nikon Plan Apo Lambda; 0.75 NA, 1 mm WD; MRD00205) with the following settings: GFP excitation 488 nm, emission 510-540 nm, exposure 500 ms, EM gain 150, digitizer 10 MHz (14-bit); ATTO647N excitation 650 nm, emission 665-705 nm, exposure 800 ms, EM gain 300, digitizer 10 MHz (14-bit); BF DIC, exposure 500 ms, EM gain 200, digitizer 10 MHz (14-bit). After imaging, fish were euthanized with an overdose of anaesthetic (0.068% MS-222).

Prior to imaging of zebrafish juveniles, individuals were anaesthetized in 0.017% MS-222, after which they were positioned on a pre-warmed petri dish containing 1% agarose within grooves moulded to the shape of a juvenile zebrafish to aid imaging and covered with 0.017% MS-222. Zebrafish juveniles were imaged with a Leica M205FA fluorescence stereo microscope (Leica Microsystems), using a 25x magnification with the following settings: BF excitation, exposure 8 ms, gain 1; GFP excitation, 800 ms, gain 1.2; ATTO647N (CY5) excitation, exposure 10.6 s, gain 3. After imaging, fish were euthanized with an overdose of anaesthetic (0.068% MS-222).

### Statistical analysis

Statistics were performed using GraphPad Prism (version 10.2.1). The specific statistical tests used are indicated in the figure legends. For all statistics, p<0.05 was considered significant.

## Results

### A mass ratio of 6:1 capsid protein to plasmid is optimal to encapsulate pVax-GFP

Prior to delivery into fish cells *in vitro* and zebrafish larvae and juveniles *in vivo*, we assessed the optimal capsid protein (CPs) to plasmid ratio that could encapsulate pVax-GFP. As indicated by the shift in DNA migration on an agarose gel, at ratios between 2:1 and 5:1, CPs bind to pVax-GFP but not all pVax-GFP is encapsulated (Fig 1). At a ratio of 6:1 or higher however, complete encapsulation was observed. Similarly, also for encapsulation of *GFP-mRNA* a 6:1 ratio was selected as previously reported [6,20].

**Fig 1.**
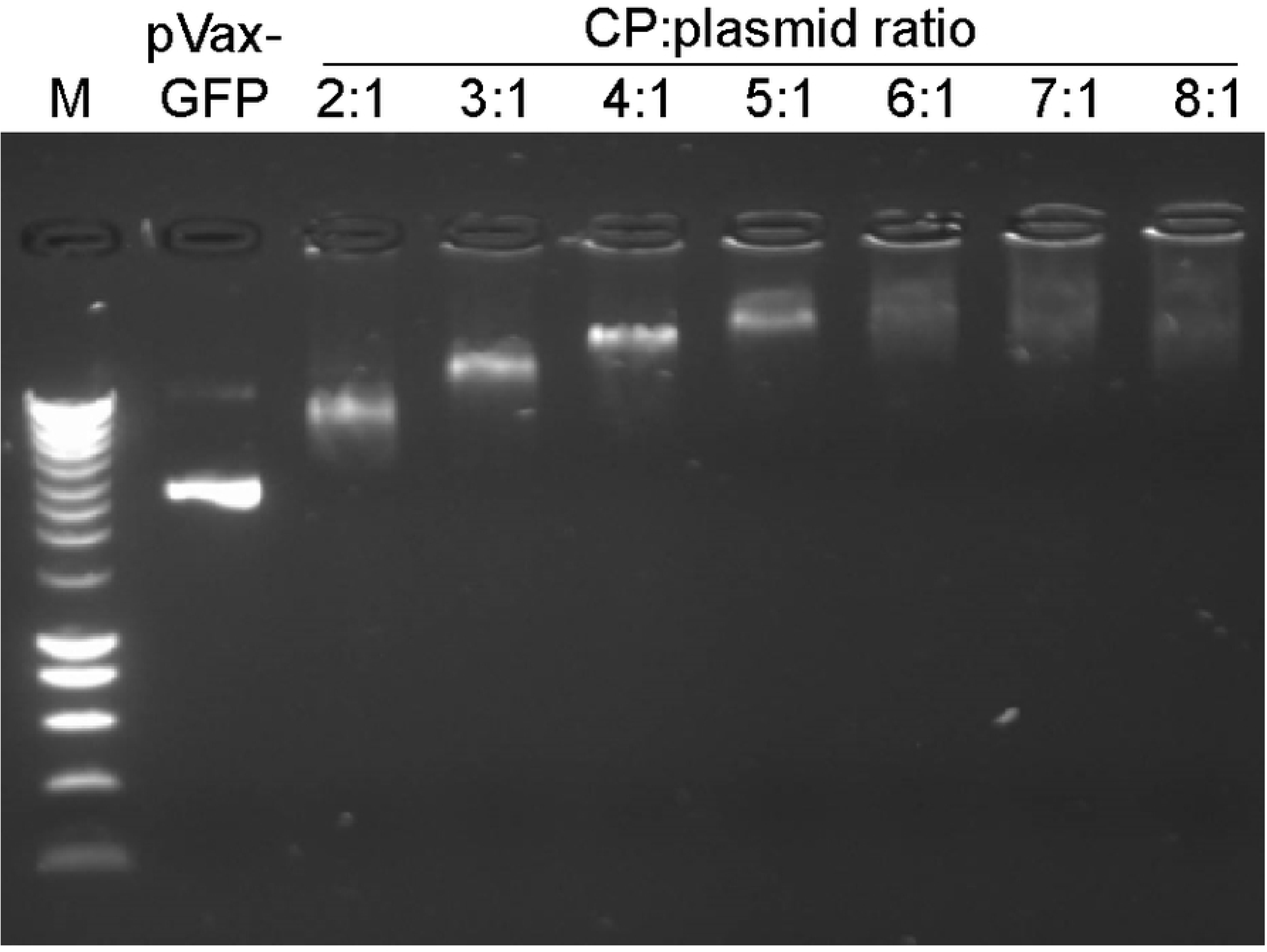
A mass ratio of 6:1 CP:plasmid is optimal to encapsulate pVax-GFP. CP and pVax-GFP (3690 bp) were mixed at the indicated ratios in assembly buffer at pH 7.2, incubated overnight at 4°C, after which the products were loaded and run on a 1% agarose gel. Free pVax-GFP was used as a control (Lane 2). M: 200 to 10000 bp molecular weight marker.

### Stability of VLPs depends on pH and cargo

The VLPs were next assessed for stability in buffers at different pH. To this end, CPs and pVax-GFP were mixed in assembly buffer at pH 7.2 and either stored in the same buffer (VLPs-DNA pH 7.2) or dialysed against VLP-DNA buffer at pH 5.0 (VLPs-DNA pH 7.2–5.0). In all conditions, we observed the formation of tubular VLPs of different lengths (Fig 2A). While at pH 7.2 only tubular VLPs were observed, at pH 5.0 spherical VLPs could also be seen (Fig 2A, right panel), likely due to the assembly of excess CPs into empty VLPs. Independent of pH, VLPs-DNA were stable for up to 5 weeks.

**Fig 2.**
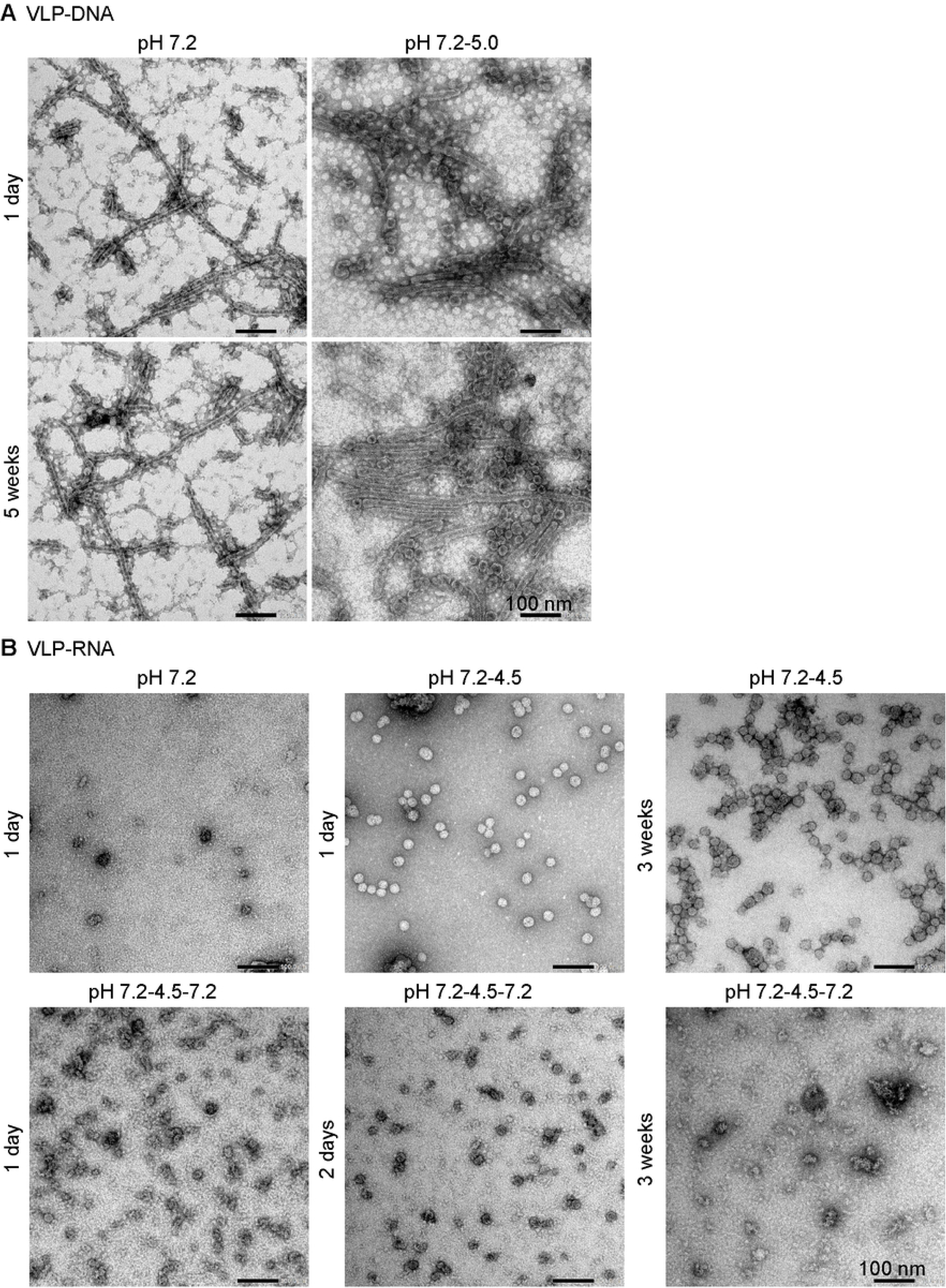
Stability of VLPs depends on the pH and cargo. VLPs were stained with 2% uranyl acetate and analysed by transmission electron microscopy. **A)** CPs were mixed at a 6:1 ratio (w/w) with pVax-GFP in assembly buffer at pH 7.2, after which part of the VLPs-DNA pH 7.2 were dialysed against VLP-DNA buffer at pH 5.0. **B)** CPs were mixed at a 6:1 ratio (w/w) with *GFP-mRNA* in assembly buffer at pH 7.2, after which part of the VLPs-RNA pH 7.2 were dialysed to VLP-RNA buffer at pH 4.5; a proportion of these was then dialysed again against assembly buffer at pH 7.2.

To assess the morphology and stability of VLPs-RNA, CPs and *GFP-mRNA* were mixed in assembly buffer at pH 7.2 (VLPs-RNA pH 7.2); a portion was then dialysed against VLP-RNA buffer at pH 4.5 and either stored in the same buffer (VLPs-RNA pH 7.2–4.5) or dialysed again against assembly buffer at pH 7.2 (VLPs-RNA pH 7.2–4.5– 7.2). Spherical VLPs-RNA, stable for up to 3 weeks, were observed only when VLPs-RNA were stored in buffer at pH 4.5 (Fig 2B, upper panel). As soon as these were brought to pH 7.2, they disassembled within 24 hours; after 6 days, only aggregates were observed (Fig 2B, lower panel).

### When added directly to the culture medium, VLPs can enter fish cells but cannot transfer nucleic acids

Next, using fluorescently labelled ATTO647N-VLPs, we evaluated whether VLPs could be taken up by EPC and HeLa cells; HeLa cells acted as a positive control as they were previously shown to take up VLPs [7]. Both, EPC cells (Fig 3A) and HeLa cells (S1A Fig) were able to take up ATTO647N-VLPs-DNA and ATTO647N-VLPs-RNA, as confirmed by the orthogonal view analysis (Fig 3B; S1B Fig). Then we examined whether VLPs could transfer nucleic acids to the cells *in vitro* using GFP expression as reporter for nucleic acids transfer. To this end, EPC and HeLa cell lines were exposed to VLPs-DNA pH 7.2, VLPs-DNA pH 7.2–5.0, VLPs-RNA pH 7.2–4.5 and VLPs-RNA pH 7.2–4.5–7.2; the latter VLPs and HeLa cells were included because it was previously shown that VLPs-RNA pH 7.2–4.5–7.2 were able to transfer *GFP-mRNA* to HEK293, HeLa and HK2 cells *in vitro* [6]. The other VLPs were selected based on their stability in their respective buffers (Fig 2). After 3 days, GFP expression was assessed by fluorescent microscopy. No GFP-positive cells were observed in EPC cells (Fig 3C) and HeLa cells (S1C Fig), suggesting that VLPs cannot transfer nucleic acids to fish cells for subsequent reporter gene expression.

**Fig 3.**
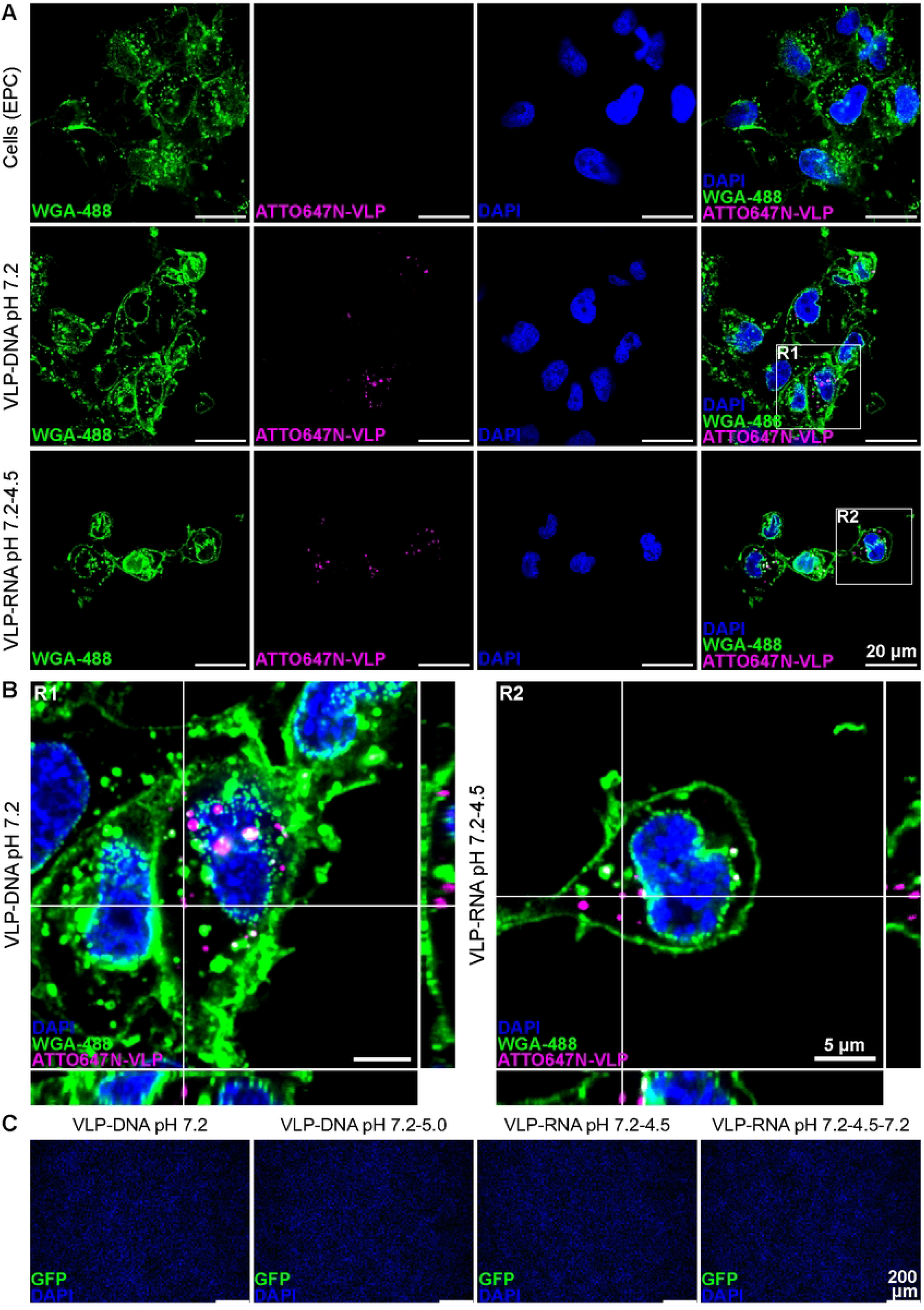
VLPs can enter but cannot transfer nucleic acids to EPC cells. VLPs were assembled by mixing CPs or ATTO647N-labelled CPs with pVax-GFP or GFP-mRNA in a 6:1 ratio. **A-B)** ATTO647N-VLPs-DNA pH 7.2 equivalent to 0.8 µg pVax-GFP, and ATTO647N-VLPs-RNA pH 7.2-4.5 equivalent to 0.5 µg GFP-mRNA were added to EPC cells, and incubated for 4 hours at 27°C. Next, cells were fixed with 4% PFA, the nuclei were stained with DAPI and the cell surface was stained with WGA-488 (5 µg/mL). **A)** Images were taken using the Nikon Ti Eclipse PFS3 microscope in combination with the Andor Revolution spinning disk confocal microscope at 100x magnification. **B)** Orthogonal view from 2 selected regions (R1+R2). **C)** VLPs-DNA at the indicated pH and equivalent to 0.8 µg pVax-GFP, or VLPs-RNA at the indicated pH and equivalent to 0.5 µg GFP-mRNA were added to EPC cells, and incubated for 3 days at 27°C. Cells cultured with VLPs-DNA were fixed with 4% PFA, and the nuclei were stained with DAPI. Cells cultured with VLPs-RNA were fixed with 4% PFA, permeabilized with triton X-100, and then stained with 1 µg/mL Chicken (IgY) polyclonal anti-GFP, followed by 2 µg/mL Goat Anti-Chicken IgY Alexa Fluor-488. The nuclei were stained with DAPI, and images were acquired with a DMi8 fluorescence microscope.

### VLPs can transfer nucleic acids to fish cells after transfection

After having observed that VLPs could be taken up by EPC cells but could not transfer nucleic acids into the cells, we next assessed whether VLPs could transfer nucleic acids when transfected into EPC cells. Transfection generally circumvents the phagocytic and endocytic entry route into cells, and direct delivery of VLPs into the cytoplasm is assumed. To this end, EPC cells were exposed to VLPs-DNA pH 7.2, VLPs-DNA pH 7.2–5.0, VLPs-RNA pH 7.2, VLPs-RNA pH 7.2–4.5 and VLPs-RNA pH 7.2–4.5–7.2, which were nuclease-treated (or not). In EPC cells transfected with VLPs-DNA pH 7.2, GFP-positive cells were observed, and at a similar percentage compared to cells transfected with naked pVax-GFP. Furthermore, a lower percentage of GFP-positive cells was observed when cells were transfected with VLPs-DNA pH 7.2–5.0 compared to VLPs-DNA pH 7.2 (Fig 4A and 4B). VLPs-DNA pH 7.2 and VLPs-DNA pH 7.2–5.0 can both protect the DNA cargo against DNase treatment, as comparable fluorescence was observed between untreated and DNase-treated samples (Fig 4A and 4B), but clearly, VLPs-DNA pH 7.2–5.0 are less able than VLP-DNA pH 7.2 to transfer DNA to the cells.

**Fig 4.**
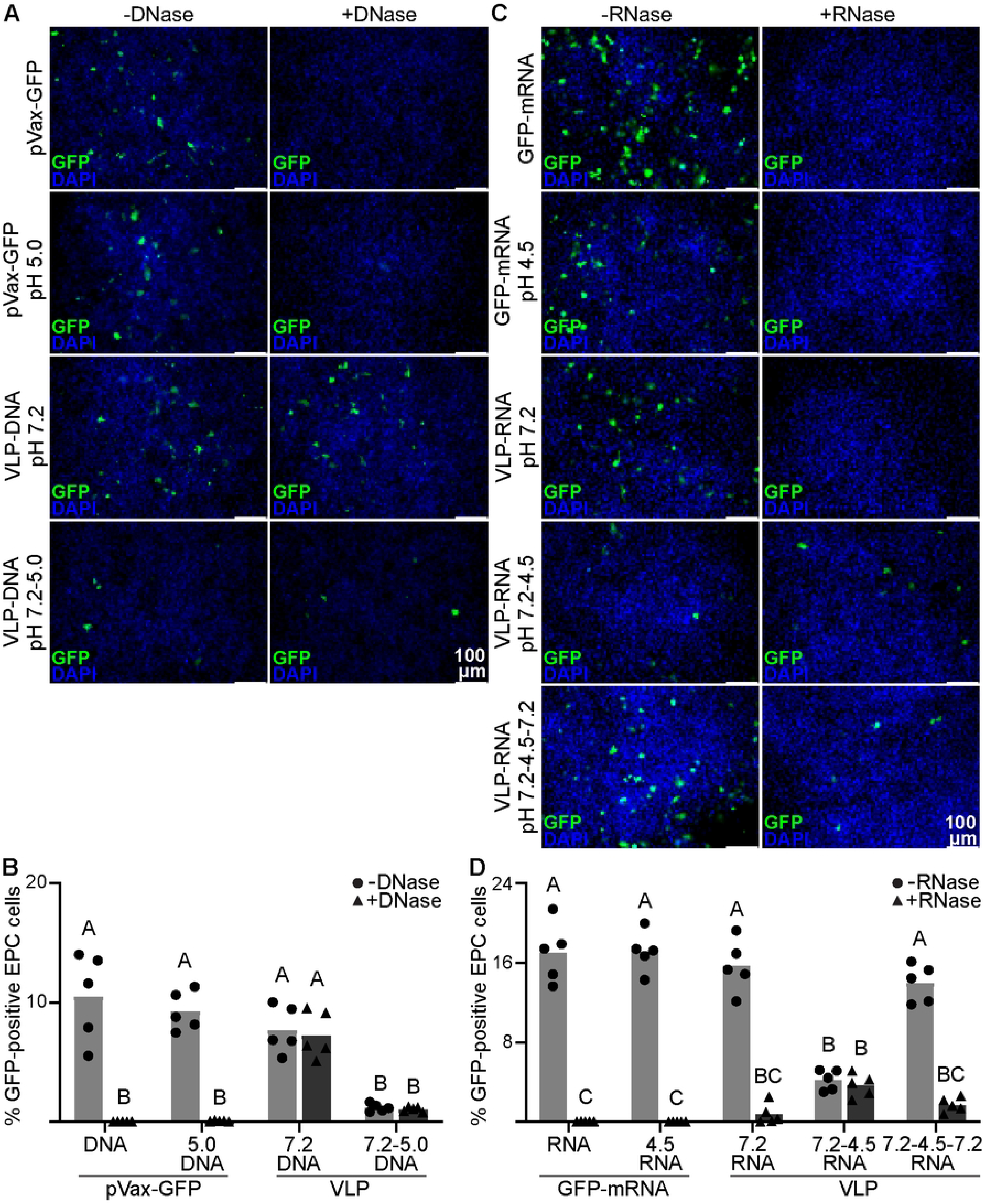
VLP-mediated nucleic acid transfer to fish cells after transfection. VLPs were assembled by mixing CPs with pVax-GFP or GFP-mRNA in a 6:1 ratio. **A and B)** EPC cells were transfected with 0.8 µg pVax-GFP, 0.8 µg pVax-GFP at pH 5.0, or VLPs-DNA at the indicated pH and equivalent to 0.8 µg pVax-GFP. DNase treatment was performed at the same pH at which the VLPs-DNA were stored and prior to mixing with the transfection reagent. **C and D)** EPC cells were transfected with 0.5 µg GFP-mRNA, 0.5 µg GFP-mRNA at pH 4.5, or VLPs-RNA at the indicated pH and equivalent to 0.5 µg GFP-mRNA. RNase treatment was performed at the same pH at which the VLPs-RNA were stored and prior to mixing the samples with the transfection reagent. **A, C)** After 3 days, cells transfected with pVax-GFP or VLPs-DNA were fixed with 4% PFA and the nuclei were stained with DAPI; whereas for cells transfected with *GFP-mRNA* or VLPs-RNA, cells were fixed with 4% PFA, permeabilized with triton X-100, and then stained with 1 µg/mL Chicken (IgY) polyclonal anti-GFP, followed by 2 µg/mL Goat Anti-Chicken IgY Alexa Fluor-488. The nuclei were stained with DAPI, and images were acquired with a DMi8 fluorescence microscope. Shown are images from one representative experiment out of five performed independently. **B, D)** After 3 days, cells transfected with pVax-GFP or VLPs-DNA were collected and resuspended in FACS buffer; cells transfected with *GFP-mRNA* or VLPs-RNA were collected and permeabilized/fixed with BD Cytofix/Cytoperm, after which were stained with 1 µg/mL Chicken (IgY) polyclonal anti-GFP, followed by 2 µg/mL Goat Anti-Chicken IgY Alexa Fluor-488. The percentage of GFP-positive cells was quantified by flow cytometry. Bars represent the average of n=5 independent experiments. Differences were analysed by two-way ANOVA with the Geisser–Greenhouse correction (assuming no sphericity) followed by Šídák’s multiple comparisons test.

In EPC cells transfected with VLPs-RNA, VLPs-RNA pH 7.2, VLPs-RNA pH 7.2–4.5 and VLPs-RNA pH 7.2–4.5–7.2 were all able to deliver nucleic acids to the cells, but VLPs-RNA pH 7.2–4.5 were less efficient than the other VLPs-RNA, as assessed by the lower percentage of GFP-positive cells (Fig 4C and 4D). Despite the lower percentage of GFP-positive cells, VLPs-RNA pH 7.2–4.5, but not VLPs-RNA pH 7.2 and VLPs-RNA pH 7.2–4.5–7.2, were able to protect RNA from degradation. The latter results are in agreement with our results from the transmission electron microscopy analysis (Fig 2B), showing that while VLPs-RNA pH 7.2–4.5 retain a spherical shape encapsulating RNA, VLP-RNA pH 7.2 consisted of capsid protein/RNA aggregates and VLPs-RNA pH 7.2–4.5–7.2 disassembled when dialysed from pH 4.5 back to pH 7.2 (Fig 2B).

### Inhibition of endosomal acidification does not lead to the release of nucleic acids by VLPs

After having shown that VLPs can transfer nucleic acids to cells when delivered to the cytoplasm by transfection, and that the efficiency was lower when VLPs were stored at pH 4.5-5.0 instead of pH 7.2, we hypothesized that the uptake mechanism, likely via the endosomes [7], in combination with enhanced stability of VLPs at low pH (≤5.0), might be the reason why VLPs cannot transfer nucleic acids to the cells. In this scenario, the acidic environment of the endosomes would stabilize the CPs-nucleic acids interaction to such an extent that it would hinder the release of nucleic acids into cells. To test this hypothesis, we investigated whether inhibition of endosomal acidification using chloroquine would promote VLPs disassembly. Chloroquine is known to be able to increase the pH of the endolysosome [42,43]. Prior to this, the optimal concentration and toxicity of chloroquine was determined for use in EPC and HeLa cells (S2 Fig). Next, EPC and HeLa cells were pre-treated for 5-10 minutes with 2, 10 and 50 µM chloroquine, after which the cells were exposed to VLPs-DNA pH 7.2, VLPs-DNA pH 7.2-5.0, VLPs-RNA pH 7.2–4.5 and VLPs-RNA pH 7.2–4.5–7.2 and tested for GFP reporter expression; in none of the conditions tested VLPs were able to transfer nucleic acids and lead to subsequence GFP expression in EPC cells (S3 Fig) nor HeLa cells (S4 Fig).

### dfTAT enhances VLPs-DNA uptake in EPC and HeLa cells

Having demonstrated that inhibition of endosomal acidification did not lead to nucleic acid release from VLPs, we next tested whether reducing the time VLPs spend in the endosome would result in nucleic acid release, as visualized by GFP reporter gene expression. To this end, we used the cell-penetrating peptide dfTAT (dimer fluorescent TAT), dfTAT originates from the HIV-1 TAT (Trans-Activator of Transcription) protein, and is able to cause endosomal leakage, leading to direct cytosolic delivery [44,45]. We hypothesize that when taken up concomitantly with VLPs, dfTAT would cause endosomal leakage, allowing VLPs to subsequently escape the endosome. To this end, first, we tested whether dfTAT was taken up by EPC cells. HeLa cells were included as control as dfTAT was already shown to cause endosomal leakage, leading to direct cytosolic delivery in these cells [44,45]. EPC and HeLa cells were exposed to 5 µM dfTAT, ATTO647N-VLPs-DNA pH 7.2, or to ATTO647N-VLPs-DNA pH 7.2 in combination with 5 µM dfTAT. In EPC and HeLa cells dfTAT was taken up within 2 hours in EPC cells (Fig 5A; S5 Fig), and 3 to 4 hours in HeLa cells (Fig 5A; S6 Fig). After 3 to 5 hours, no difference in dfTAT uptake was observed between HeLa cells incubated with dfTAT alone or with dfTAT in combination with ATTO647N-VLPs-DNA pH 7.2; in EPC cells however, a higher dfTAT uptake was observed in cells exposed to dfTAT in combination with ATTO647N-VLPs-DNA pH 7.2 compared to cells exposed to dfTAT alone (Fig 5A; S5 Fig). A difference in ATTO647N-VLPs-DNA pH 7.2 uptake was observed in EPC and HeLa cells incubated with ATTO647N-VLPs-DNA pH 7.2 alone, compared to EPC and HeLa cells incubated with ATTO647N-VLPs-DNA pH 7.2 in combination with dfTAT. After 5 hours of incubation, the median ATTO647 fluorescent intensity in HeLa and EPC cells incubated with ATTO647N-VLPs-DNA pH 7.2 in combination with dfTAT was significantly higher than in cells incubated with VLPs-DNA pH 7.2 alone, with the largest difference (from 1297 to 91134) observed in EPC cells (Fig 5B; S5 and S6 Figs).

**Fig 5.**
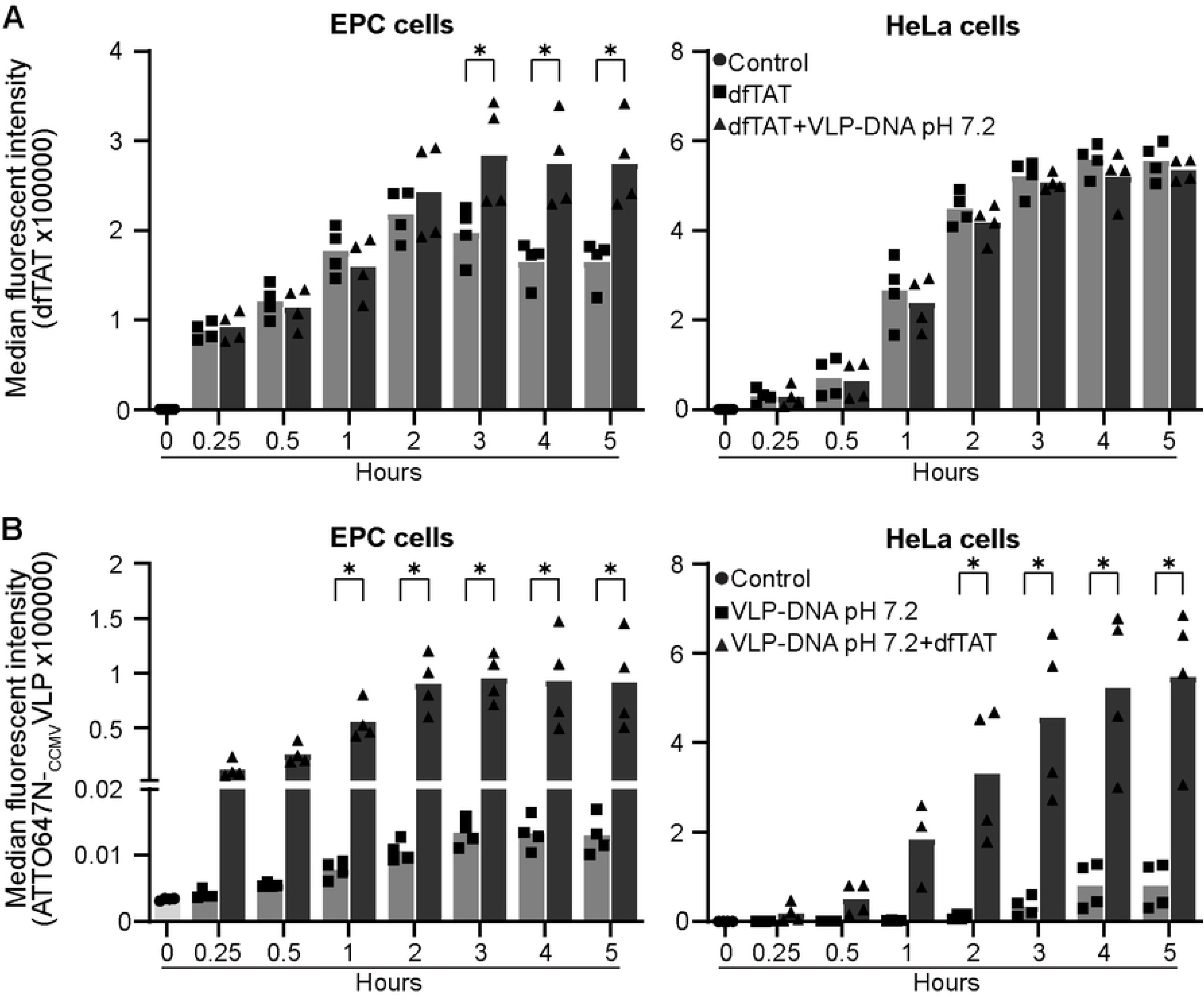
dfTAT enhances VLPs-DNA uptake in EPC and HeLa cells. VLPs were assembled by mixing ATTO647N-labelled CPs and pVax-GFP in a 6:1 ratio. ATTO647N-VLPs-DNA pH 7.2, equivalent to 0.8 µg pVax-GFP, 5 µM dfTAT, or a combination of ATTO647N-VLPs-DNA pH 7.2 with 5 µM dfTAT, were added to EPC or HeLa cells, and incubated for 0 to 5 hours. Cells were washed with PBS, trypsinized and resuspended in FACS buffer; fluorescence was measured by flow cytometry. Bar graphs represent the average of n=4 independent experiments. Differences were analysed by two-way ANOVA with the Geisser–Greenhouse correction (assuming no sphericity) followed by Šídák’s multiple comparisons test. **A)** dfTAT fluorescence. **B)** ATTO647N-VLPs-DNA pH 7.2 fluorescence.

### dfTAT promotes DNA transfer by VLPs-DNA to HeLa cells

Having shown the uptake of dfTAT by EPC and HeLa cells, and that this leads to substantially more VLPs-DNA pH 7.2 uptake than in this absence of dfTAT, we next tested whether the presence of dfTAT also promotes the transfer of nucleic acids. To this end, EPC and HeLa cells were exposed to VLPs-DNA pH 7.2, VLPs-DNA pH 7.2-5.0 and VLPs-RNA pH 7.2–4.5 in combination with 5 µM dfTAT. These VLPs were selected based on their stability in their respective buffers (Fig 2) and their ability to protect nucleic acid cargo (Fig 4). However, in none of the conditions tested were VLPs able to transfer nucleic acids and lead to reporter gene expression (Fig 6A-C; S7 Fig), except in HeLa cells exposed to VLPs-DNA pH 7.2 in combination with 5 µM dfTAT (Fig 6A-C). To rule out the possibility that dfTAT was used at sub-optimal concentrations, HeLa cells were exposed to VLPs-DNA pH 7.2 in combination with increasing concentrations of dfTAT (1-40 µM). A dose-dependent increase in GFP-positive cells was seen in cells receiving 1, 2.5, 5 and 10 µM dfTAT (Fig 6D), whereas cytotoxicity was observed in cells exposed to > 10 µM dfTAT (data not shown). The highest percentage of GFP-positive cells was observed in cells that received VLPs-DNA pH 7.2 in combination with 5 or 10 µM dfTAT (Fig 6D). Similar conditions were tested in EPC cells, but no GFP-positive cells were observed (S7 Fig).

**Fig 6.**
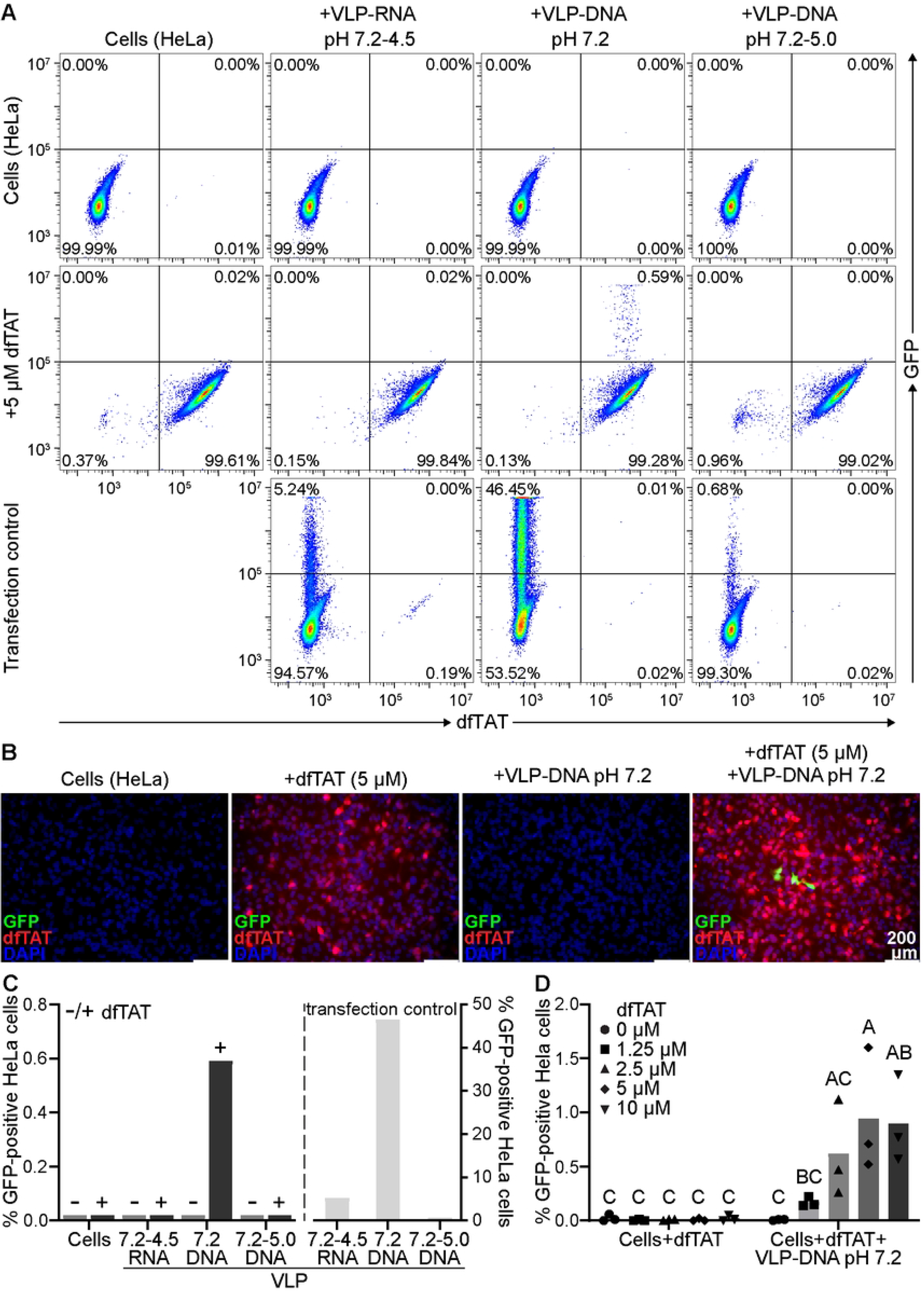
dfTAT promotes DNA transfer by VLPs-DNA to HeLa cells. VLPs were assembled by mixing CPs or ATTO647N-labelled CPs with pVax-GFP or GFP-mRNA in a 6:1 ratio. VLPs-DNA at the indicated pH and equivalent to 0.8 µg pVax-GFP, or VLPs-RNA at the indicated pH and equivalent to 0.5 µg GFP-mRNA, in combination with 5 µM dfTAT were added to HeLa cells, and incubated for 5 hours at 37 °C, after which the medium was replaced for culture medium. **A and C)** After 2 days, cells exposed to VLPs-DNA were collected and resuspended in FACS buffer; cells exposed to VLPs-RNA were collected and permeabilized/fixed with BD Cytofix/Cytoperm, after which they were incubated with 1 µg/mL Chicken (IgY) polyclonal anti-GFP, followed by 2 µg/mL Goat Anti-Chicken IgY Alexa Fluor-488. The percentage of GFP-positive cells was quantified by flow cytometry. Bar graph represents one experiment. **B)** After 2 days, cells were fixed with 4% PFA, and the nuclei were stained with DAPI. Images were acquired with a DMi8 fluorescence microscope. **D)** VLPs-DNA pH 7.2 equivalent to 0.8 µg pVax-GFP in combination with 1.25, 2.5, 5 and 10 µM dfTAT were added to HeLa cells and incubated for 5 hours at 37 °C, after which the medium was replaced with culture medium. After 2 days, cells were collected, and the percentage of dfTAT-positive and GFP-positive cells was quantified by flow cytometry. Bar graph represents the average of n=3 independent experiments. Differences were analysed by two-way ANOVA with the Geisser–Greenhouse correction (assuming no sphericity) followed by Šídák’s multiple comparisons test.

### VLPs-DNA pH 7.2 are able to transfer DNA *in vivo* to zebrafish upon intramuscular injection

After having demonstrated VLPs uptake into EPC and HeLa cells *in vitro*, but absence of nucleic acids delivery, with the exception of HeLa cells exposed to VLPs-DNA pH 7.2 in combination with the cell-penetrating dfTAT, we next tested whether VLPs could be taken up and transfer DNA *in vivo* in zebrafish larvae and juveniles. The advantage of an *in vivo* model is the presence of more cell types that could possibly take up VLPs, thereby better reflecting the natural situation. To this end, optically transparent casper zebrafish larvae were intramuscularly injected with DNase-treated pVax-GFP (negative control), pVax-GFP (positive control), and DNase-treated ATTO647N-VLPs-DNA pH 7.2. DNase treatment was included to ensure that GFP expression was due to the delivery of pVax-GFP by the VLPs-DNA pH 7.2, and not due to residual non-encapsulated pVax-GFP. VLPs-DNA pH 7.2 were selected because of their stability (Fig 2) and ability to protect the DNA cargo in earlier experiments (Fig 4). While ATTO647N fluorescence could be observed in all zebrafish larvae injected with ATTO647N-VLPs-DNA pH 7.2 (S1 Video), GFP-positive myocytes, resulting from GFP reporter expression, were observed in n=8 of the n=35 injected zebrafish (Fig 7A, S2 Video). After pVax-GFP injection, the positive control, GFP-positive myocytes were observed in n=7 out of n=35 zebrafish (Fig 7A, S2 Video), indicating a similar GFP reporter expression in zebrafish intramuscularly injected with naked pVax-GFP DNA or ATTO647N-VLPs DNA pH 7.2. Although macrophages and neutrophils recruited to the injection site were observed to take up ATTO647N-VLPs-DNA pH 7.2 (S8 Fig), they did not seem to contribute to the observed GFP expression patterns. In fact, besides GFP-positive myocytes, no other single GFP-positive cells, consistent with the location of macrophages or neutrophils, were observed (Fig 7A, S2 Video). To further substantiate these findings, zebrafish juveniles (5-week-old) were intramuscularly injected with pVax-GFP and DNase-treated ATTO647N-VLPs-DNA pH 7.2. Whereas autofluorescence was observed in the gut area of all fish (bottom left corner of each image; Fig 7B), GFP-positive myocytes were observed in both groups with similar efficiencies (Fig 7B), indicating successful DNA delivery by ATTO647N-VLPs-DNA pH 7.2 also into zebrafish juveniles after intramuscular injection.

**Fig 7.**
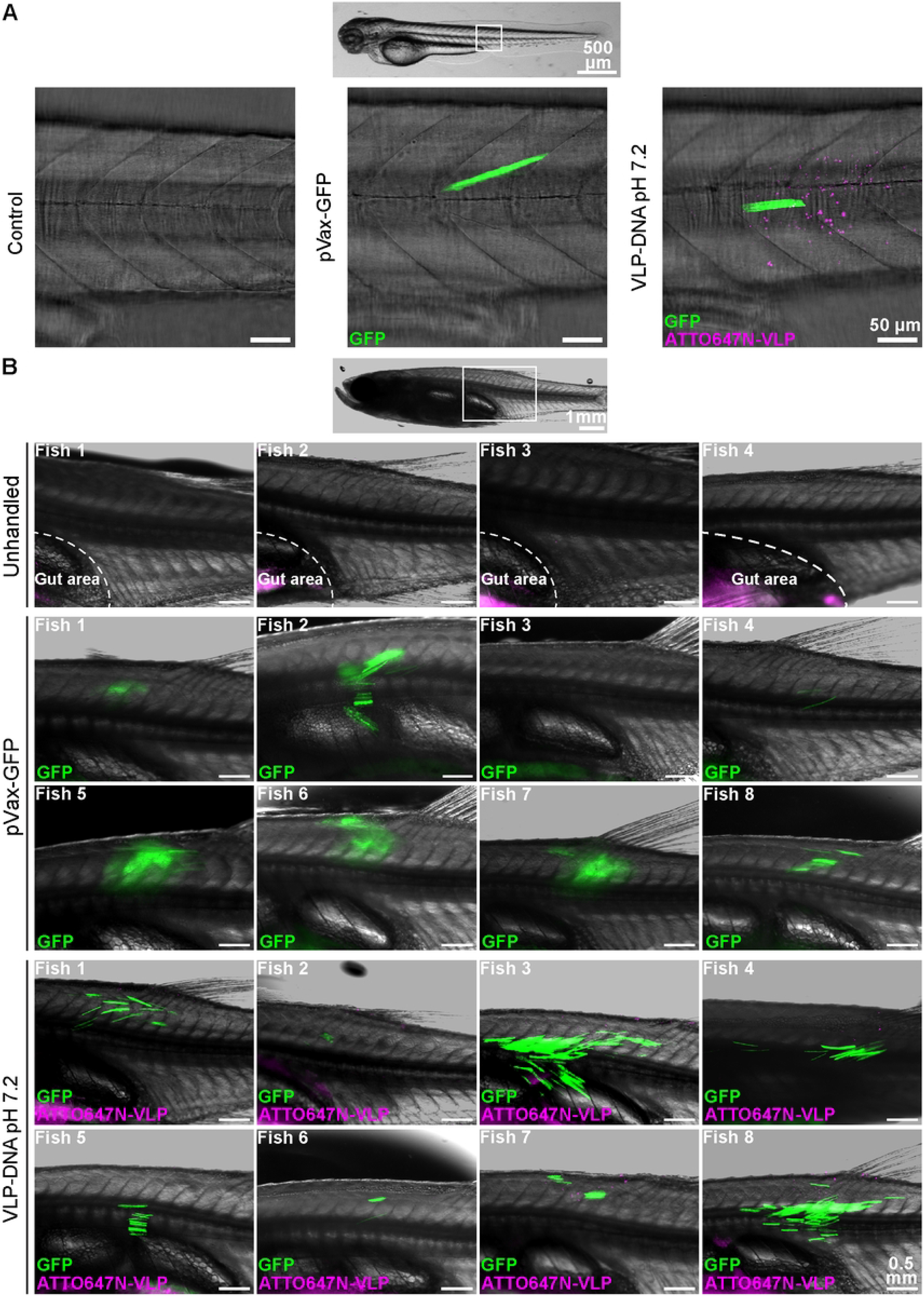
ATTO647N-VLPs-DNA pH 7.2 mediated transfer of DNA to zebrafish larvae and juveniles. ATTO647N-VLPs were assembled by mixing ATTO647N-labelled CPs with pVax-GFP in a 6:1 ratio. **A)** Casper zebrafish larvae (3 dpf) were injected intramuscularly with 2 nl of DNase-treated 0.207 mg/mL pVax-GFP (control), 2 nl of 0.207 mg/mL pVax-GFP or 2 nl of DNase-treated 0.207 mg/mL ATTO647N-VLPs-DNA pH 7.2. Zebrafish were imaged 5 days post-injection using a Nikon Ti Eclipse PFS3 microscope in combination with the Andor Revolution spinning disk confocal. Shown are images from one representative fish. The entire stack, including the stacks of the biological replicates (from the fish with GFP-positive myocytes), are provided as supplementary video 2. **B)** Casper zebrafish juveniles (5-week-old) were either unhandled or injected intramuscularly with 1-2 µL of 0.221 mg/mL pVax-GFP or 1-2 µL of DNase-treated 0.221 mg/mL ATTO647N-VLPs-DNA pH 7.2. Zebrafish were imaged 8 days post-injection using a Leica M205FA fluorescence stereo microscope.

## Discussion

In this study, the potential of CCMV VLPs to encapsulate and transfer nucleic acids to animal cells *in vitro,* and to zebrafish larvae and juveniles *in vivo* was investigated. The results showed that, independent of the pH of the storage buffer, VLPs-DNA were stable for up to 5 weeks, whereas VLPs-RNA were stable for up to 3 weeks but only when stored in a buffer at pH 4.5. When administered directly to the culture medium, VLPs were taken up by EPC and HeLa cells but could not unpack and transfer nucleic acids to the cells, whereas VLPs delivered by transfection could. When VLPs were administered directly to the culture medium along with the cell-penetrating peptide dfTAT, only VLPs-DNA pH 7.2 were able to transfer the DNA to HeLa but not EPC cells. *In vivo,* VLPs-DNA pH 7.2 could transfer DNA to myocytes in zebrafish larvae and juveniles upon intramuscular injection.

In this study, VLPs were assembled containing *GFP-mRNA* (∼1168 nt) or pVax-GFP (3690 bp). Most studies used CP to nucleic acid ratios between 4.1-6.5 to 1 to encapsulate nucleic acids of various lengths [4–8,20–22,24,46–48]. For *GFP-mRNA* encapsulation, a 6:1 CP to RNA (w/w) ratio was selected based on previous literature. For pVax-GFP encapsulation, the optimal CP to DNA ratio was optimized during this study because previous reports were limited, particularly for plasmids of similar size to pVax-GFP.

Transmission electron microscopy showed that VLPs-RNA were stable for up to 3 weeks but only when VLPs-RNA were stored in buffer at pH 4.5. As soon as these were brought to pH 7.2, they disassembled within 24 hours. This observation was in line with our data on the delivery of RNA to fish cells after transfection in which only VLPs-RNA pH 7.2-4.5 were able to protect RNA against RNase treatment, whereas VLPs-RNA pH 7.2-4.5-7.2 were not. It is known that when VLPs and RNA are mixed at pH 7.2, the CP-CP interactions are too weak to assemble into (stable) virus particles. When the pH is lowered however, the CP-CP interactions are strengthened, ultimately leading to complete capsids with a spherical morphology similar to that of the wild-type virus [5,49]. While we observed that VLPs-RNA pH 7.2-4.5-7.2 disassembled within 24 hours when the pH was brought back to pH 7.2, Garmann observed that VLPs-RNA pH 7.2-4.5-7.2 were stable for at least 1 week at 4°C [5]. There are two differences between the assembly protocols. The first, is that Garmann mixed CP and RNA in protein buffer (20 mM Tris-HCl, 1 M NaCl, 1 mM EDTA, 1 mM DTT and 1 mM PMSF), after which the CP-RNA mixture was dialysed against assembly buffer, whereas our CP and RNA were directly mixed in assembly buffer. The second difference is the ionic strength of the buffer: in our study, VLPs-RNA were generated and stored in an assembly buffer containing 50 mM Tris-HCl, 100 mM NaCl, 10 mM KCl, and 5 mM MgCl2, whereas Garmann used the same buffer but with 50 mM instead of 100 mM NaCl. These relatively small differences in ionic strength, however, do impact the interactions between the negatively charged cargo and the positive interior of the CP. In that way, provisionally, explaining the observed differences in VLPs-RNA pH 7.2-4.5-7.2 stability.

Furthermore, our results showed that VLPs-RNA pH 7.2-4.5 were stable and could protect the RNA against RNase treatment. Similar results were reported by Azizgolshani, who showed that VLPs-RNA pH 7.2-4.8 were able to protect the RNA cargo against RNase treatment [21]. In contrast, Karan reported substantial RNA degradation (∼90%) following RNase treatment on VLPs-RNA pH 7.2-4.5, explained by the assembly of VLPs with longer RNA transcripts (4433 and 4626 nt) [4]. In fact, the encapsulation of such long RNA transcripts is often shared among multiple VLP capsids (multiplets), leaving a portion of the RNA molecule accessible and prone to nuclease degradation [20]. In contrast to VLPs-RNA, VLPs-DNA were stable already at pH 7.2. A previous study showed that mixing CP with DNA fragments of different lengths (100, 200, 500, and 1000 bp) in buffer at pH 7.2 leads to the formation of rod-shaped VLPs-DNA [7]. However, whether rod-shaped VLPs-DNA protect DNA against DNase treatment is not known. Our study shows that both VLPs-DNA pH 7.2 and VLPs-DNA pH 7.2-5.0 are stable for up to five weeks and protect the DNA against DNase treatment.

We demonstrated that VLPs could transfer nucleic acids to cells when delivered to the cytoplasm by transfection. It is unlikely that this was the result of partial or complete disassembly of VLPs after interaction with the transfection mix. In fact, when cells were transfected with VLPs-RNA pH 7.2-4.5, we consistently observed a lower percentage of GFP-positive cells compared to cells transfected with VLPs-RNA pH 7.2-4.5-7.2 or naked RNA, indicating that the RNA was still encapsulated within VLPs-RNA pH 7.2-4.5 despite the presence of the transfection reagent. The lower percentage of GFP-positive cells is likely due to the higher stability of the VLP-RNA interaction at pH 4.5 compared to pH 7.2, as further discussed later.

Contradictory results have been reported on whether VLPs can unpack and transfer nucleic acids to animal cells *in vitro*. Similar to our study, it was reported that VLPs-RNA pH 7.2-4.8 containing *eYFP-RNA* were able to unpack and transfer RNA to BHK-21 cells only after transfection with Lipofectamine 2000, whereas BHK-21 cells to which VLPs were directly added to the cell culture medium did not show eYFP expression [21]. Furthermore, the same study observed a lower efficiency of RNA transfer in cells transfected with VLPs-RNA pH 7.2-4.8 than in cells transfected with naked *eYFP-RNA*. This is consistent with observations in our study in which the efficiency of nucleic acid transfer was lower when VLPs pH 4.5-5.0 were transfected than when naked nucleic acids were transfected. Another study confirmed that Lipofectamine is required for successful nucleic acid delivery to HeLa cells when using VLPs that have encapsulated mRNA-DNA origami encoding GFP [50]. Conversely, in another study, VLPs-RNA pH7.2-4.5-7.2 encapsulating *eGFP-mRNA* were able to transfer mRNA to HEK293, HeLa and HK2 cells when added directly to the cell culture medium, without using a transfection reagent [6]. The latter study used VLPs-RNA pH 7.2-4.5-7.2 that were dialysed against assembly buffer pH 7.2 for a minimum of 4 hours. In our study, we used VLPs-RNA pH 7.2-4.5-7.2 dialysed against assembly buffer pH 7.2 for 24 hours. Since both studies used the same buffers, the only difference could be the time that VLPs-RNA pH 7.2-4.5-7.2 were at pH 7.2. Unfortunately, the study by Villagrana-escareño did not specify for how long the VLPs-RNA pH 7.2-4.5-7.2 were at pH 7.2 prior to administration to the cells, nor did they include an RNase treatment [6]. In our study, we observed that when VLPs-RNA were brought from pH 4.5 back to pH 7.2, they disassembled within 24 hours, as assessed by electron microscopy analysis and by the fact that they did not protect the RNA against RNase treatment.

In general, efficient delivery of nucleic acids by VLPs into animal cells *in vitro* remains challenging. To gain further insights, it is important to understand the natural transmission cycle of CCMV. CCMV is transmitted when beetles, primarily the bean leaf beetle (*Cerotoma trifurcate*) and spotted cucumber beetle (*Diabrotica undecimpunctata*), feed on infected host plants (cowpea, soybean, mung bean or common bean) [51,52]. During feeding, CCMV particles from infected plants are ingested by the beetle and then enter the digestive tract. The pH within the digestive tract is slightly acidic, favouring the stability of the CCMV capsid. CCMV does not replicate within the beetle; viral particles are retained in the gut and are transmitted during subsequent feeding on healthy plants. When the infected beetle feeds on a new plant, the viral particles are delivered directly into the plant cell’s cytoplasm. Within the cytoplasm, at a pH of approximately 7.0-7.5, the capsid swells and forms pores through which the viral RNA is released directly into the cytoplasm, after which the positive-sense RNA genome can be directly translated by host ribosomes [53–56]. The CCMV genome encodes a movement protein that helps the virus spread from an infected cell to neighbouring cells via plasmodesmata, which are cytoplasmic channels connecting plant cells [53,57].

The biggest difference between the natural transmission cycle of CCMV and its use as a VLP in animal cells is that, in the natural situation, CCMV is directly delivered to the cytoplasm of a plant cell (pH 7.0-7.5), whereas in animal cells, CCMV is taken up in endosomes (pH <6.5). Previous studies reported that VLPs-DNA and VLPs-RNA are taken up in the endosomes of HeLa cells [7,22]. Within endosomes, the pH drops from ∼6.5 in early endosomes to ∼5.5 in late endosomes and ∼4.5 in lysosomes [43]. Combining our results showing that efficiency of nucleic acid transfer to EPC cells is lower when VLPs pH 4.5-5.0 were transfected compared to when VLPs pH 7.2 were used, with reports showing that VLPs are taken up in endosomes (pH <6.5) and that VLPs have enhanced stability at pH ≤6, we hypothesized that the acidic environment of the endosomes stabilizes the VLPs to such an extent that prevents the release of nucleic acids into cells. Inhibition of endosomal acidification however, did not lead to the release of nucleic acids by VLPs. There could be two reasons for this: first, the increase in endosomal pH is not sufficient to cause swelling of the VLPs for nucleic acids release. In mouse peritoneal macrophages, 100 µM chloroquine increased lysosomal pH from 4.7 to 6.3 within 2-3 minutes. When chloroquine was removed, the effect was partially reversed within minutes [58]. Second, nucleic acids are released from the VLPs but remain packed within the endosome. Studies with RNA have shown that ∼99% of endocytosed RNA fails to enter the cytoplasm. Taking into account that RNA is partially degraded in the endosome and cytoplasm, it was estimated that ∼0.3% of endocytosed RNA is present in the cytoplasm [59].

Comparing the natural situation, in which CCMV delivery into the cytoplasm of a plant cell (pH 7.0-7.5) directly leads to a co-translational disassembly, to the situation in animal cells, in which VLPs could transfer nucleic acids to HeLa and EPC cells only when directly delivered to the cytoplasm by transfection, we assessed a strategy to deliver VLPs into the cytoplasm using the cell-penetrating peptide dfTAT. Despite the presence of dfTAT, VLPs-RNA pH 7.2–4.5 were not able to transfer RNA to cells and only VLPs-DNA pH 7.2 were able to transfer DNA to HeLa cells, although the efficiency was ∼79x lower than when VLPs were transfected, likely due to the very high stability of VLPs-DNA pH 7.2. Yet, the mechanism by which VLPs-DNA pH 7.2 transfer the DNA to the cells remains unclear. When considering VLPs-RNA however, it is possible that the low pH (pH 4.5) step has stabilized the CPs-nucleic acids interaction to such an extent that it is likely the particles were degraded in the endolysosome before they could release the RNA into the cytoplasm.

Finally, when DNase-treated VLPs-DNA pH 7.2 were delivered *in vivo* by intramuscular injection, they were able to unpack and transfer DNA to myocytes of zebrafish larvae and juveniles. It is possible that due to the pressure at which particles are administered upon intramuscular injection, VLPs-DNA pH 7.2 are not first internalised via endosomes, but are rather directly delivered to the cytoplasm, resembling the situation in which the cell-penetrating peptide was present *in vitro* or when a beetle feeds on a plant *in vivo*. Only a limited number of studies have investigated the use of VLPs to deliver nucleic acids *in vivo*. Upon subcutaneous injection of VLPs encapsulating replicon RNA encoding GFP into the footpads of mice, GFP expression was observed in the draining lymph node [4]. In the latter study however, while *in vitro* they reported ∼90% degradation upon RNase treatment, explained by the assembly of 4433 nt RNA in multiplets that do not fully protect the RNA, *in vivo* they injected non-RNase treated VLPs. Other studies successfully used encapsulated oxidized DNA (ox-dsDNA), CpG ODN 1826 and Poly(I:C), both *in vitro* and *in vivo* [8,46,48]. A main difference is that upon endosomal uptake, oxDNA and Poly(I:C) can be recognized by endosomal Toll-like receptors [60,61], while in our system, the nucleic acids require endosomal escape for subsequent protein expression in the cytoplasm.

In conclusion, CCMV VLPs enable a straightforward and effective encapsulation method for nucleic acid cargos, providing protection against nuclease degradation, with minor differences in stability depending on the type of nucleic acid and pH storage conditions. Despite the fact that CCMV can be produced at high yields in plants, with relatively low costs, making it suitable for scale-up for vaccine production purposes, based on our findings, the key limitation of the CCMV VLPs system is that upon direct *in vitro* delivery to animal cells, CCMV VLPs are taken up in the acidic environment of the endosomes. This, stabilises the VLPs rather than support nucleic acids’ release to the cells for subsequent protein expression. The addition of a cell-penetrating peptide, which leads to direct delivery of VLPs-DNA pH 7.2 into the cytoplasm, had limited effect on nucleic acids transfer efficiency. *In vivo* however, intramuscular administration of VLPs-DNA pH 7.2 resulted in the successful transfer of DNA to myocytes in zebrafish larvae and juveniles. Altogether, these findings provide the foundations to apply VLPs as delivery vehicle in fish; however, future efforts should include strategies that enable direct cytosolic delivery, to increase the chance of plasmid transfer to host cells, as well as VLPs delivery via the mucosal route (orally or by immersion) to avoid injection of individual animals.

## Acknowledgments

The authors would like to thank Junika Pohl, Amy Broos and Ines Lobato Dionisio (Wageningen University), whose work laid the foundation for the CCMV research; Kaylee van Dijk for her assistance with zebrafish injections and spinning-disk microscopy.

## Supporting information

**S1 Fig.**
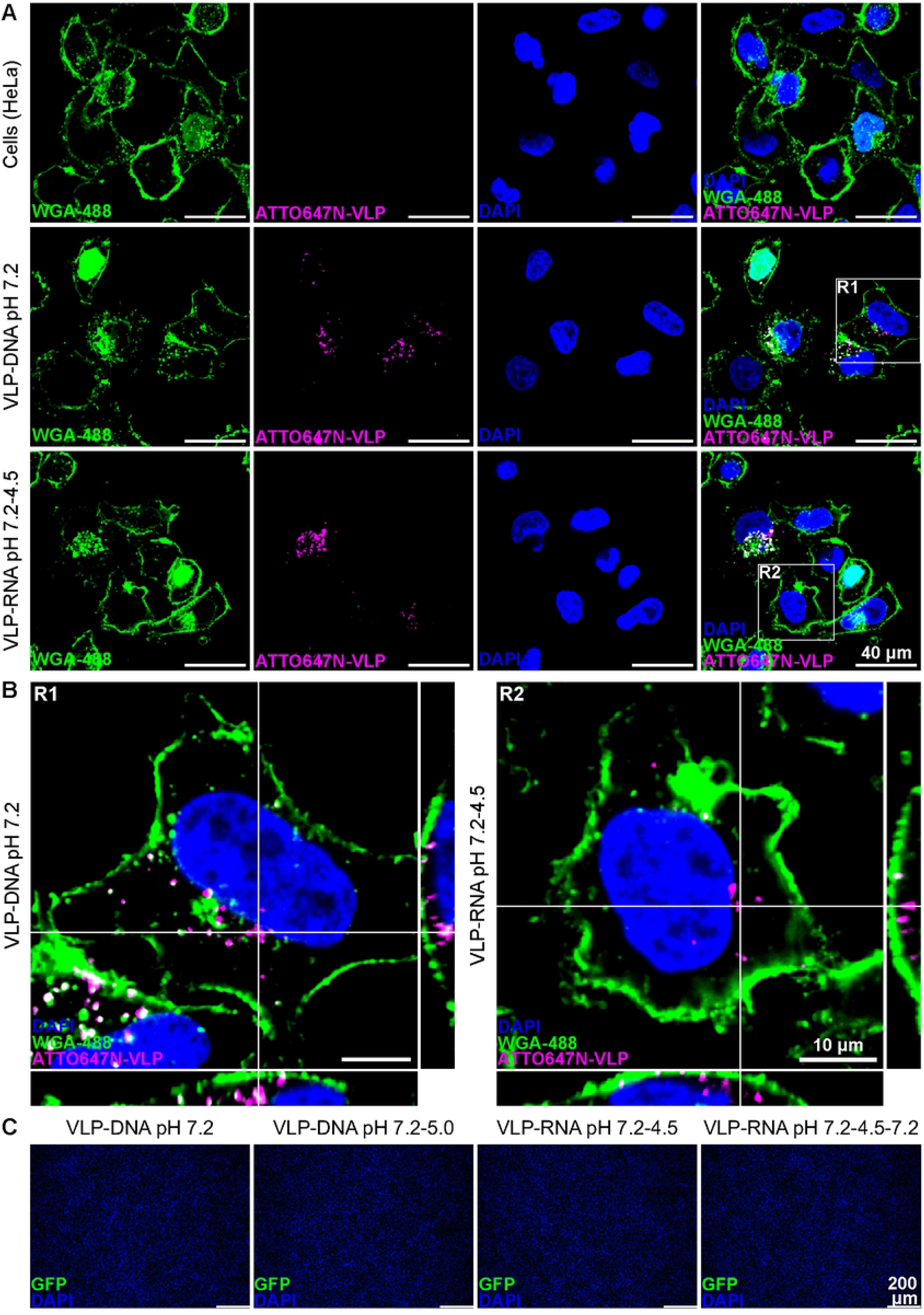
VLPs can enter HeLa cells but cannot transfer nucleic acids. VLPs were assembled by mixing CPs or ATTO647N-labelled CPs with pVax-GFP or GFP-mRNA in a 6:1 ratio. **A and B)** ATTO647N-VLPs-DNA pH 7.2 equivalent to 0.8 µg pVax-GFP, and ATTO647N-VLPs-RNA pH 7.2-4.5 equivalent to 0.5 µg GFP-mRNA were added to HeLa cells, and incubated for 4 hours at 27°C. Next, cells were fixed with 4% PFA, the nuclei were stained with DAPI and the cell surface was stained with WGA-488 (5 µg/mL). **A)** Images were taken using the Nikon Ti Eclipse PFS3 microscope in combination with the Andor Revolution spinning disk confocal microscope at 60x magnification. **B)** Orthogonal view from 2 selected regions (R1+R2). **C)** VLPs-DNA at the indicated pH and equivalent to 0.8 µg pVax-GFP, or VLPs-RNA at the indicated pH and equivalent to 0.5 µg GFP-mRNA were added to HeLa cells, and incubated for 3 days at 37°C. Cells cultured with VLPs-DNA were fixed with 4% PFA, and the nuclei were stained with DAPI. Cells cultured with VLPs-RNA were fixed with 4% PFA, permeabilized with triton X-100, and then stained with 1 µg/mL Chicken (IgY) polyclonal anti-GFP, followed by 2 µg/mL Goat Anti-Chicken IgY Alexa Fluor-488. The nuclei were stained with DAPI, and images were acquired with a DMi8 fluorescence microscope.

**S2 Fig.**
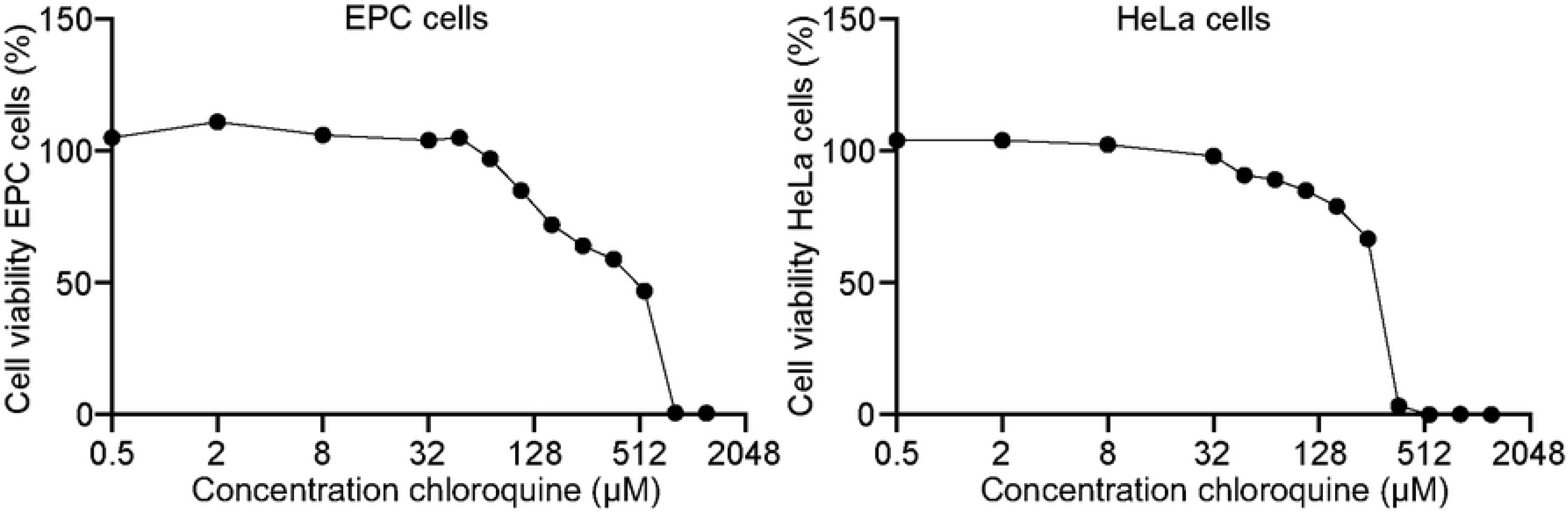
High concentrations of chloroquine are toxic for EPC and HeLa cells. EPC and HeLa cells were incubated with different concentrations of chloroquine. After 24 hours, cells were fixed with 4% PFA, stained with 1% crystal violet, after which the OD_595_ was measured. Cell viability was calculated relative to untreated samples.

**S3 Fig.**
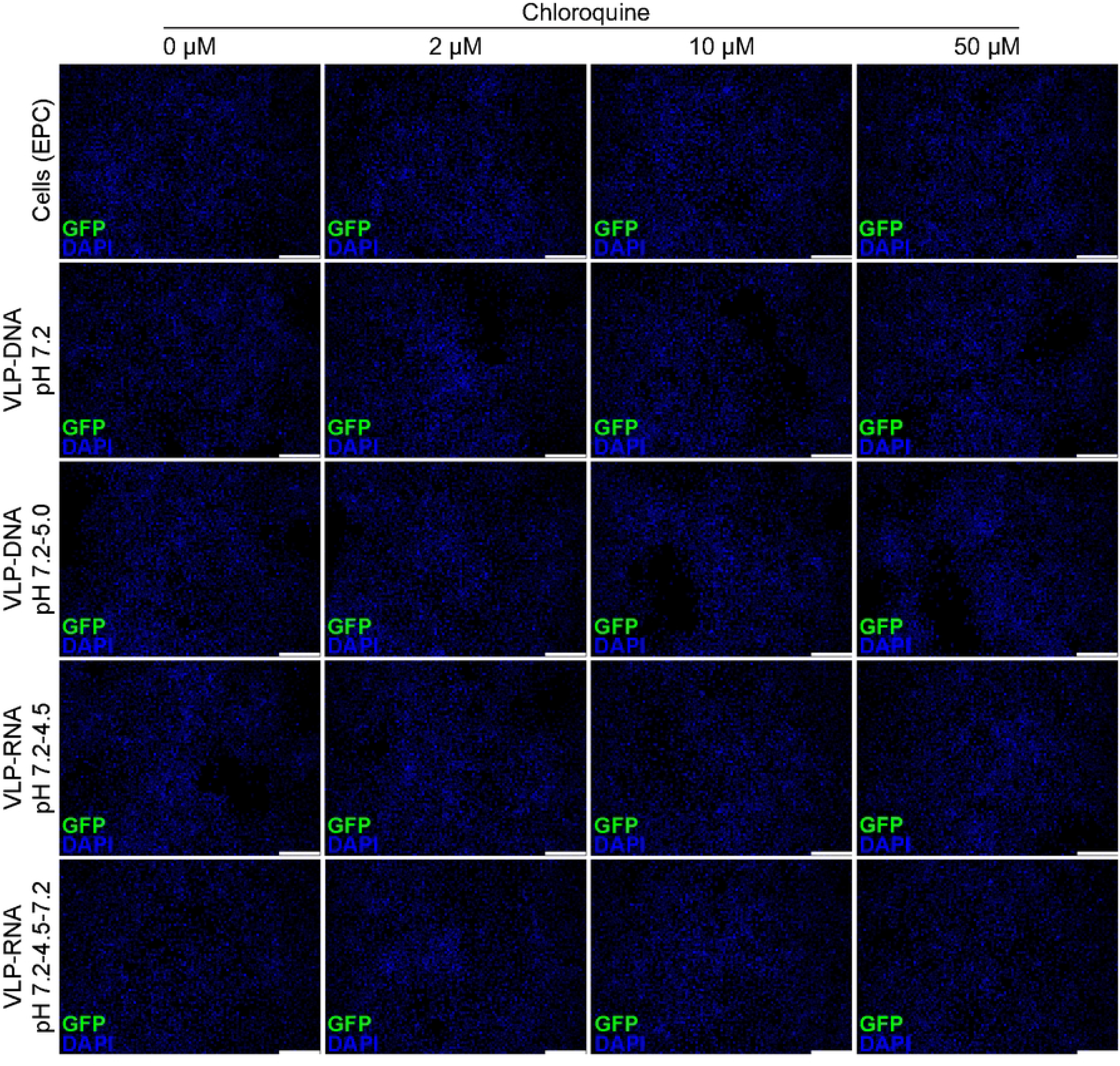
Inhibition of endosomal acidification by chloroquine does not lead to nucleic acid release by VLPs in EPC cells. VLPs were assembled by mixing CPs or ATTO647N-labelled CPs with pVax-GFP or GFP-mRNA in a 6:1 ratio. EPC cells were pre-treated with 2, 10 and 50 µM chloroquine for 10 minutes, after which VLPs-DNA at the indicated pH and equivalent to 0.8 µg pVax-GFP, or VLPs-RNA at the indicated pH and equivalent to 0.5 µg GFP-mRNA were added to the cells and incubated for 3 days at 27 °C. After 3 days, cells cultured with VLPs-DNA were fixed with 4% PFA and the nuclei were stained with DAPI. Cells cultured with VLPs-RNA were fixed with 4% PFA, permeabilized with triton X-100, and then stained with 1 µg/mL Chicken (IgY) polyclonal anti-GFP, followed by 2 µg/mL Goat Anti-Chicken IgY Alexa Fluor-488. The nuclei were stained with DAPI and images were acquired with a DMi8 fluorescence microscope.

**S4 Fig.**
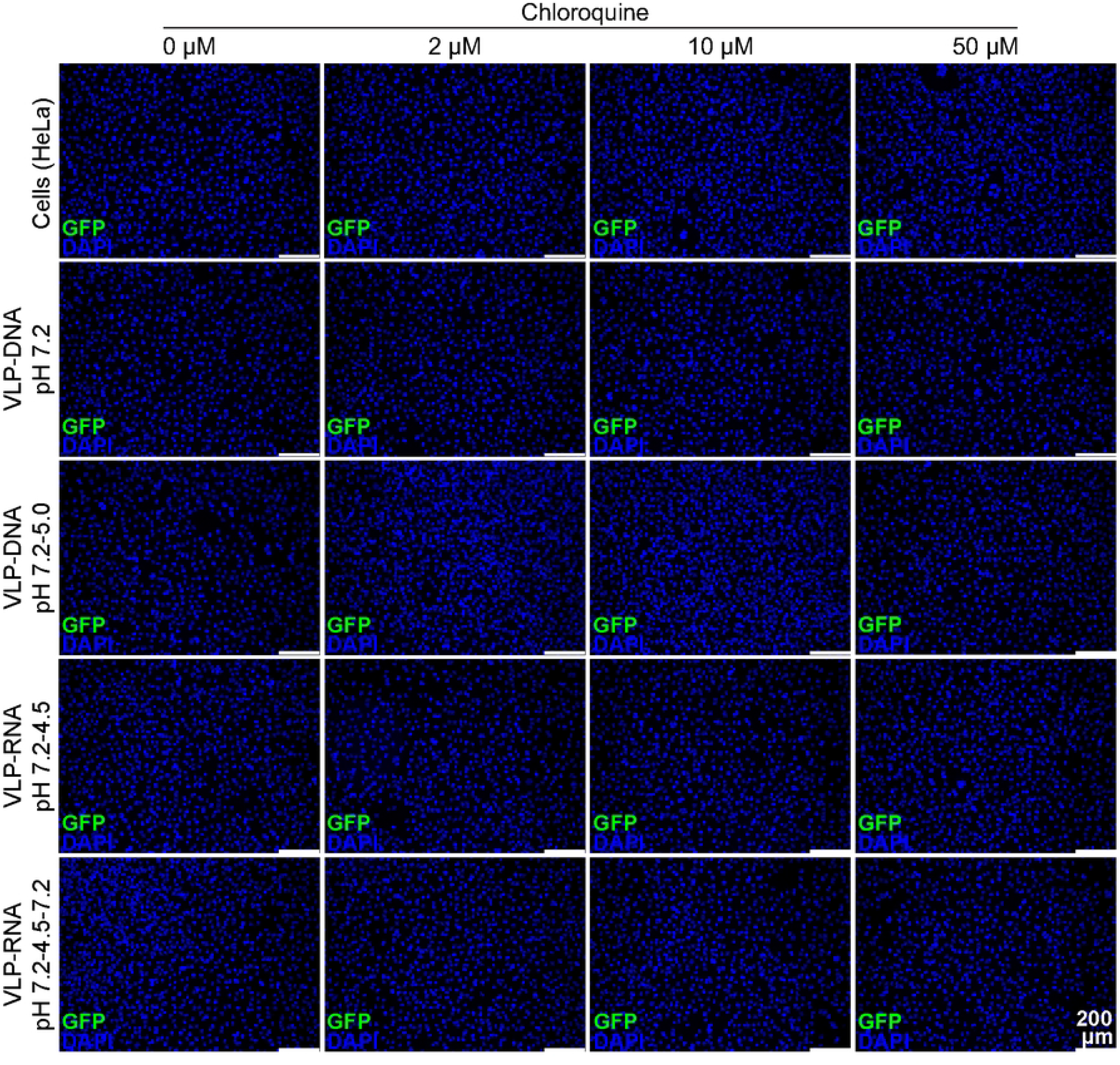
Inhibition of endosomal acidification by chloroquine does not lead to nucleic acid release by VLPs in HeLa cells. VLPs were assembled by mixing CPs or ATTO647N-labelled CPs with pVax-GFP or GFP-mRNA in a 6:1 ratio. HeLa cells were pre-treated with 2, 10 and 50 µM chloroquine for 10 minutes, after which VLPs-DNA at the indicated pH and equivalent to 0.8 µg pVax-GFP, or VLPs-RNA at the indicated pH and equivalent to 0.5 µg GFP-mRNA were added to the cells and incubated for 3 days at 37 °C. After 3 days, cells cultured with VLPs-DNA were fixed with 4% PFA and the nuclei were stained with DAPI. Cells cultured with VLPs-RNA were fixed with 4% PFA, permeabilized with triton X-100, and then stained with 1 µg/mL Chicken (IgY) polyclonal anti-GFP, followed by 2 µg/mL Goat Anti-Chicken IgY Alexa Fluor-488. The nuclei were stained with DAPI and images were acquired with a DMi8 fluorescence microscope.

**S5 Fig.**
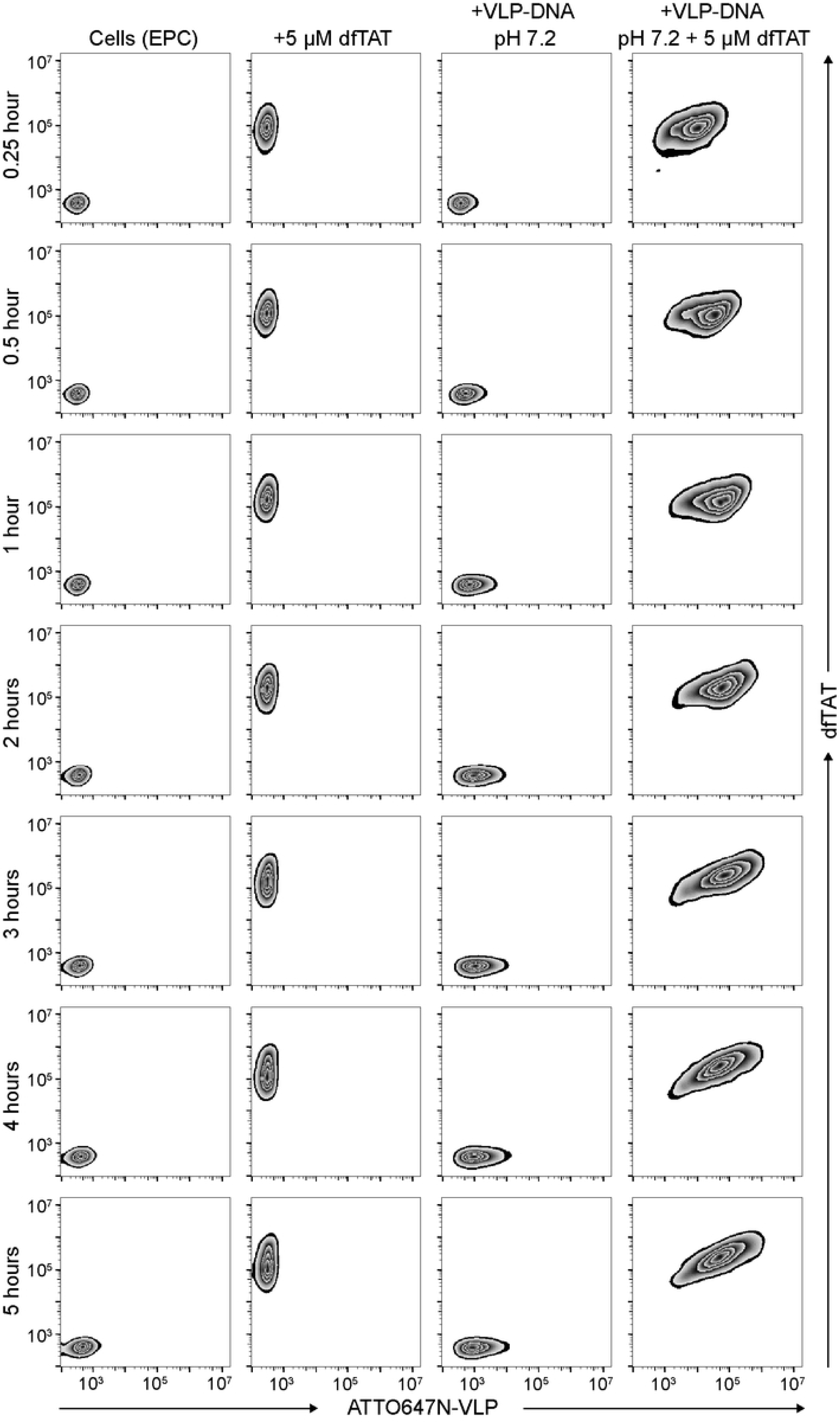
The cell-penetrating peptide dfTAT enhances VLPs-DNA uptake in EPC cells. VLPs were assembled by mixing ATTO647N-labelled CPs and pVax-GFP in a 6:1 ratio. ATTO647N-VLPs-DNA pH 7.2, equivalent to 0.8 µg pVax-GFP, 5 µM dfTAT, or a combination of ATTO647N-VLPs-DNA pH 7.2 with 5 µM dfTAT, were added to EPC cells, and incubated for 0 to 5 hours at 27 °C. Cells were washed with PBS, trypsinized and measured by flow cytometry. Flow cytometry plots correspond to the bar graphs show in Fig 5.

**S6 Fig.**
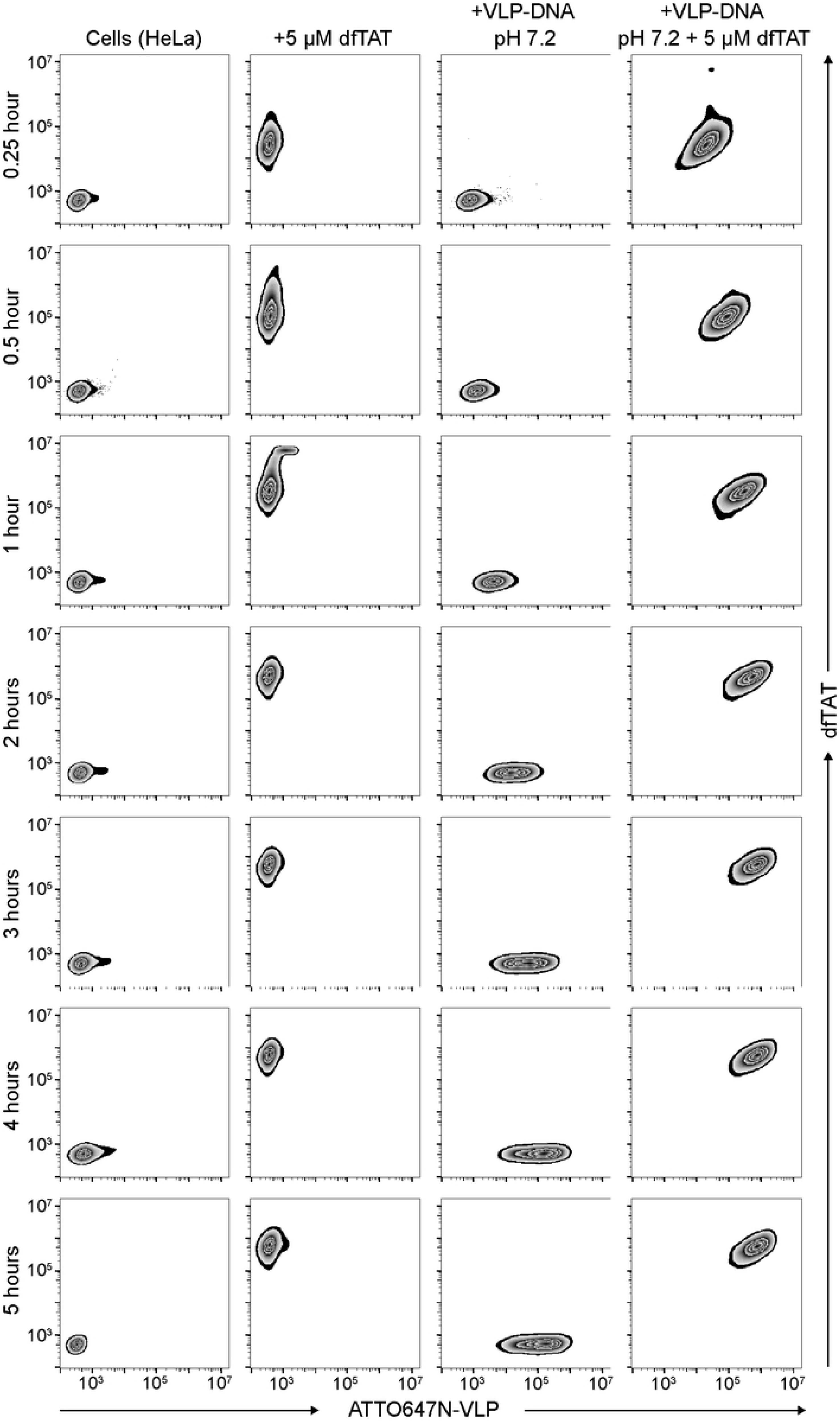
The cell-penetrating peptide dfTAT enhances VLPs-DNA uptake in HeLa cells. VLPs were assembled by mixing ATTO647N-labelled CPs and pVax-GFP in a 6:1 ratio. ATTO647N-VLPs-DNA pH 7.2, equivalent to 0.8 µg pVax-GFP, 5 µM dfTAT, or a combination of ATTO647N-VLPs-DNA pH 7.2 with 5 µM dfTAT, were added to HeLa cells, and incubated for 0 to 5 hours at 37 °C. Cells were washed with PBS, trypsinized and measured by flow cytometry. Flow cytometry plots correspond to the bar graphs show in Fig 5.

**S7 Fig.**
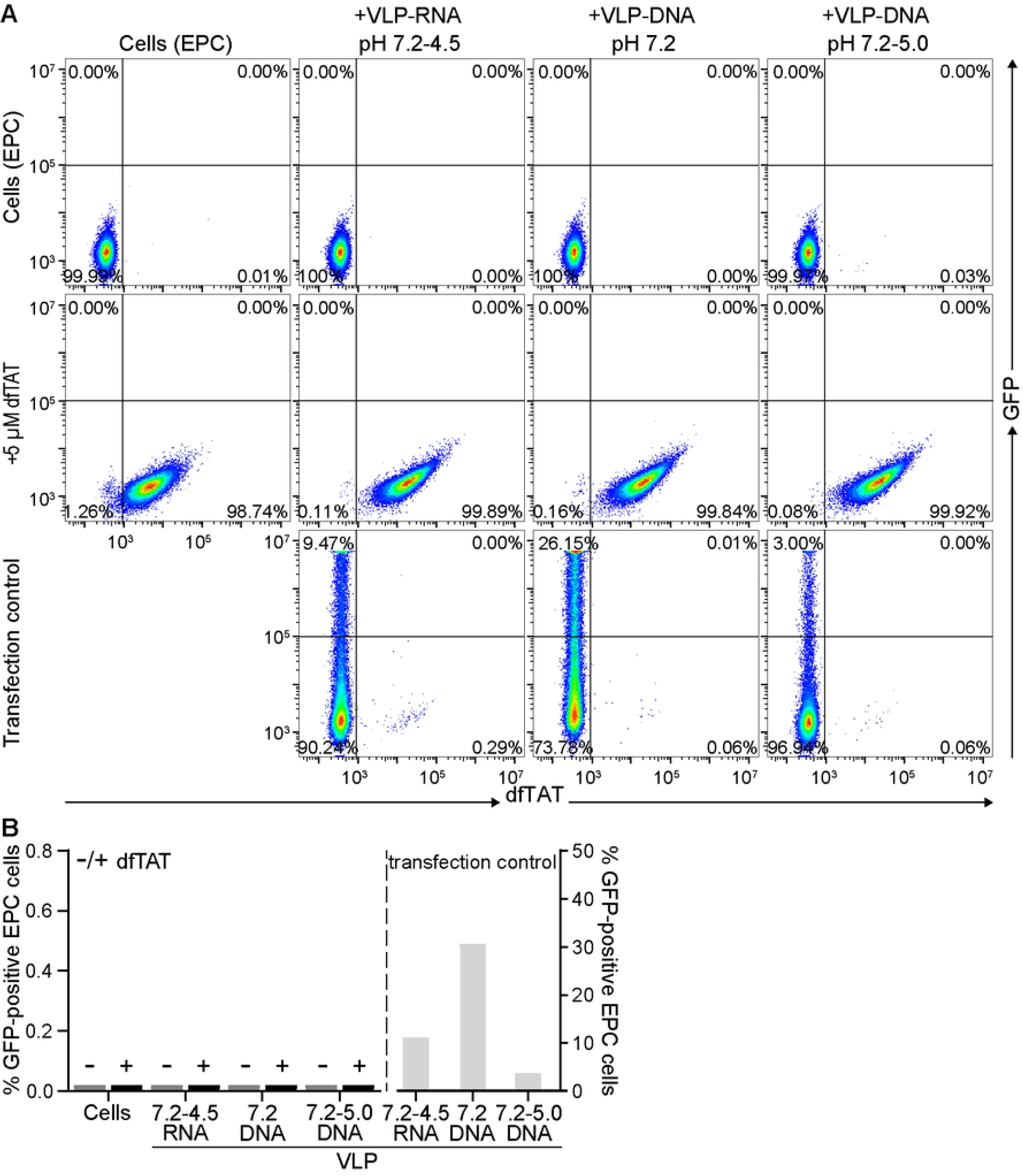
dfTAT does not enhance transfer of nucleic acids by VLPs to EPC cells. VLPs were assembled by mixing CPs or ATTO647N-labelled CPs with pVax-GFP or GFP-mRNA in a 6:1 ratio. VLPs-DNA at the indicated pH and equivalent to 0.8 µg pVax-GFP, or VLPs-RNA at the indicated pH and equivalent to 0.5 µg GFP-mRNA, in combination with 5 µM dfTAT were added to EPC cells, and incubated for 5 hours at 27 °C, after which the medium was replaced for culture medium. After 3 days, cells exposed to VLPs-DNA were collected and resuspended in FACS buffer. Cells exposed to VLPs-RNA were collected and permeabilized/fixed with BD Cytofix/Cytoperm, after which the cells were stained with 1 µg/mL Chicken (IgY) polyclonal anti-GFP, followed by 2 µg/mL Goat Anti-Chicken IgY Alexa Fluor-488. The percentage of GFP-positive cells was quantified by flow cytometry. **B)** Bar graph represents one experiment.

**S8 Fig.**
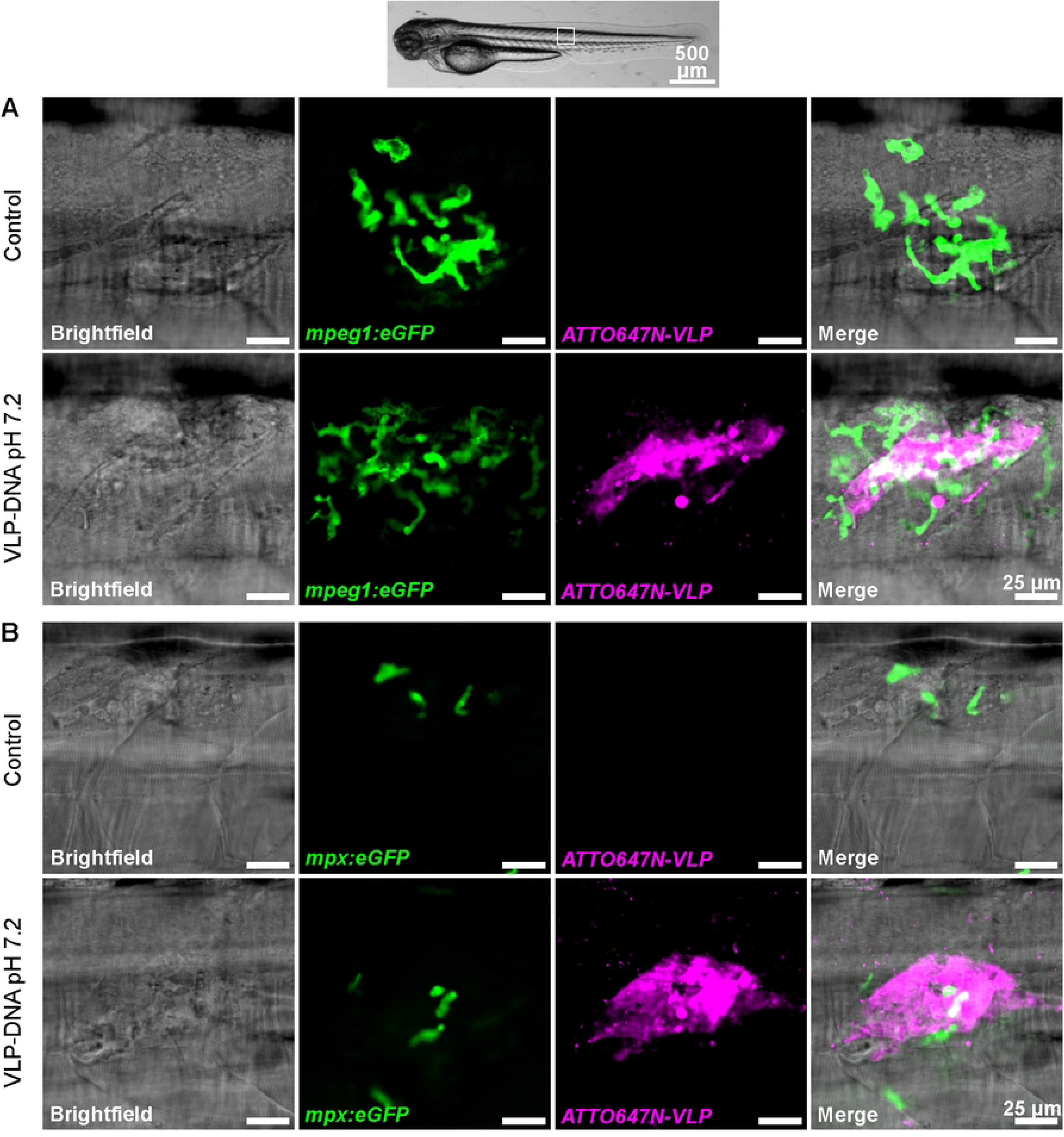
ATTO647N-VLPs-DNA pH 7.2 are taken up in macrophages and neutrophils in zebrafish larvae. ATTO647N-VLPs were assembled by mixing ATTO647N-labelled CPs with pVax-GFP in a 6:1 ratio. **A)** 3 dpf *Tg(mpeg1:eGFP)* **B)** 3 dpf *Tg(mpx-eGFP),* were injected intramuscularly with 2 nl of assembly buffer pH 7.2 (control), or 2 nl of 0.207 mg/mL ATTO647N-VLPs-DNA pH 7.2. Zebrafish were imaged 3 to 4 hours post-injection using a Nikon Ti Eclipse PFS3 microscope in combination with the Andor Revolution spinning disk confocal. Shown are images from one representative fish (out of four fish). The entire stack, including the stacks of the biological replicates, are provided as supplementary video 3 (*Tg(mpeg1:eGFP)*) and supplementary video 4 *Tg(mpx-eGFP)*.

**S1 Video. ATTO647N-VLPs-DNA pH 7.2 are present in all fish after intramuscular injection.** ATTO647N-VLPs were assembled by mixing ATTO647N-labelled CPs with pVax-GFP in a 6:1 ratio. Casper zebrafish (3 dpf, n=35 per group) were injected intramuscularly with 2 nl of DNase-treated 0.207 mg/mL pVax-GFP (control), 2 nl of 0.207 mg/mL pVax-GFP or 2 nl DNase-treated 0.207 mg/mL ATTO647N-VLPs-DNA pH 7.2. Zebrafish were imaged 5 days post-injection using a Nikon Ti Eclipse PFS3 microscope in combination with the Andor Revolution spinning disk confocal. The first n=21 zebrafish were imaged, whereas zebrafish 22 to 35 were only imaged when GFP-positive myocytes were observed.

**S2 Video. ATTO647N-VLPs-DNA pH 7.2 can transfer DNA to zebrafish larvae.** ATTO647N-VLPs were assembled by mixing ATTO647N-labelled CPs with pVax-GFP in a 6:1 ratio. Casper zebrafish (3 dpf, n=35 per group) were injected intramuscularly with 2 nl of DNase-treated 0.207 mg/mL pVax-GFP (control), 2 nl of 0.207 mg/mL pVax-GFP or 2 nl DNase-treated 0.207 mg/mL ATTO647N-VLPs-DNA pH 7.2. Zebrafish were imaged 5 days post-injection using a Nikon Ti Eclipse PFS3 microscope in combination with the Andor Revolution spinning disk confocal. In S2 Video only fish with GFP-positive myocytes are shown.

**S3 Video. ATTO647N-VLPs-DNA pH 7.2 are taken up in macrophages in zebrafish larvae.** ATTO647N-VLPs were assembled by mixing ATTO647N-labelled CPs with pVax-GFP in a 6:1 ratio. Three dpf *Tg(mpeg1-eGFP)* zebrafish were injected intramuscularly with 2 nl of assembly buffer pH 7.2 (control), or 2 nl of 0.207 mg/mL ATTO647N-VLPs-DNA pH 7.2. Zebrafish were imaged 3 to 4 hours post-injection using a Nikon Ti Eclipse PFS3 microscope in combination with the Andor Revolution spinning disk confocal. One representative fish was shown in S8 Fig, the entire stack, including the stacks of the biological replicates, are provided here as S3 Video.

**S4 Video. ATTO647N-VLPs-DNA pH 7.2 are taken up in neutrophils in zebrafish larvae.** ATTO647N-VLPs were assembled by mixing ATTO647N-labelled CPs with pVax-GFP in a 6:1 ratio. Three dpf *Tg(mpx-eGFP)* zebrafish were injected intramuscularly with 2 nl of assembly buffer pH 7.2 (control), or 2 nl of 0.207 mg/mL ATTO647N-VLPs-DNA pH 7.2. Zebrafish were imaged 3 to 4 hours post-injection using a Nikon Ti Eclipse PFS3 microscope in combination with the Andor Revolution spinning disk confocal. One representative fish was shown in S8 Fig, the entire stack, including the stacks of the biological replicates, are provided here as S4 Video.

